# Cysteine residues in the C-terminal tail of connexin32 regulate its trafficking

**DOI:** 10.1101/2020.10.18.302364

**Authors:** Anuttoma Ray, Parmender P. Mehta

**Author notes:** Correspondence should be addressed to Anuttoma Ray.

## Abstract

Gap junctions (GJ)s are formed by the assembly of constituent transmembrane proteins called connexins (Cxs). Aberrations in this assembly of Cxs are observed in several genetic diseases as well as in cancers. Hence it becomes imperative to understand the molecular mechanisms underlying such assembly defect. The polarized cells in the epithelia express Connexin32 (Cx32). The carboxyl-terminal tail (CT) of Cx32 orchestrates several aspects of GJ dynamics, function and growth. The study here was aimed at determining if post-translational modifications, specifically, palmitoylation of cysteine residues, present in the CT of Cx32, has any effect on GJ assembly. The CT of Cx32 was found to harbor three cysteine residues, which are likely to be modified by palmitoylation. The study here has revealed for the first time that Cx32 is palmitoylated at cysteine 217 (C217). However, it was found that mutating C217 to alanine affected neither the trafficking nor the ability of Cx32 to assemble into GJs. Intriguingly, it was discovered that mutating cysteine 280 and 283 in combination, blocked the transport of Cx32 from the Golgi to the cell surface. Overall, the findings reveal the importance of the two terminal cysteine residues of Cx32 in regulating its trafficking and stability and hence is ability to assemble into GJs, possibly as being part of a CAAX motif in its CT.

## INTRODUCTION

Gap junctions (GJ)s are conglomeration of several cell-cell channels at cell-to-cell contact sites involved in the direct intercellular exchange of small growth regulatory molecules (Goodenough & Paul, 2009), which permit the direct passage of molecules <1500 daltons between the cytoplasmic interiors of adjacent cells (Loewenstein, 1981). GJs are made up of proteins called connexins (Cxs). Cxs are named according to their molecular weight in kilodaltons (kDa) (Beyer et al., 1990). Study of diseases associated with mutations and knockouts (Leinonen et al.) of Cx genes in mice have provided valuable insights on Cx function and have brought forward the significance of gap junctional intercellular communication (GJIC) in maintaining tissue homeostasis (Saez et al., 2003). Defects in the expression, trafficking and assembly of Cxs into GJs have also been observed in a variety of genetic diseases (Dobrowolski & Willecke, 2009; White & Paul, 1999) as well as during progression of many carcinomas (Aasen et al., 2016; Naus & Laird, 2010) (Wu & Wang, 2019) thus highlighting the importance of GJIC in diverse physiological processes.

Connexin32 (Cx32), encoded by the *GJB1* gene, is a 32 kDa Cx, and shows a tissue-specific expression. It is present in the polarized acinar cells of exocrine glands like the pancreas, prostate, lacrimal glands, salivary glands, as well as liver hepatocytes, stomach parietal cells and oligodendrocytes (Bosco et al., 2011; Michon et al., 2005). Most studies point to a tumor-suppressive role for Cxs (Aasen et al., 2016). Studies have shown that forced expression of Cx32 in Cx-deficient human prostate cancer cell line, LNCaP, retarded tumorigenicity and induced differentiation, and these effects were contingent upon the restoration of GJ assembly (Mehta et al., 1999). In *in vivo* studies, where athymic nude mice were injected with Cx32 expressing clones of communication-deficient human tumor cells (SKHepl), a reduction in tumor growth rate was observed (Eghbali et al., 1991). The above studies implicated that tumorigeneis is negatively correlated with intercellular communication. Cx32 KO mice showed a higher incidence of spontaneous and chemically-induced liver tumors (Temme et al., 1997). An increased susceptibility to chemically-induced hepatocarcinogenesis was also observed in a transgenic mutant mouse model expressing a DN mutant (V139M) of Cx32 that caused a loss of channel function in the liver (Aasen et al., 2016). In Cx32 KO mice, X-ray irradiation also greatly enhanced liver tumorigenesis compared to the wild-type mice, as well as increased tumor formation in the lung, adrenal gland, lymph nodes and small intestine accompanied with activation of the MAPK pathway in these tumors (King & Lampe, 2004, 2004). Collectively, these studies strongly suggest that Cx32 is a tumor suppressor, at least in the context of the liver and lungs (Naus & Laird, 2010). Earlier studies showed that Cx32 is expressed in the luminal cells of the prostate whereas Cx43 in the basal cells (Habermann et al., 2002). Analysis of the distribution of Cx32 in 23 normal prostates, 43 benign prostatic hyperplasia specimens, 60 primary and 20 metastatic prostate tumors in archival and frozen sections showed that in invasive tumors, Cx32 remained intracellular whereas in well-differentiated tumors it was inefficiently assembled into GJs. Investigation into the molecular mechanisms regulating the trafficking and assembly of Cx32 into GJs in cancer cells is thus essential to enhance our understanding about the pathogenesis of different types of cancers.

The functional properties of GJ channels mainly depend on the composition of the different Cx isoforms (Bevans et al., 1998). However, channel function is also regulated by the location, type, extent, and combination of different post-translational modifications (PTMs). Cxs may undergo different types of PTMs, which includes phosphorylation, hydroxylation, acetylation, glutamation, nitrosylation, palmitoylation, and prenylation (Locke et al., 2006). The most well studied of these Cx PTMs is phosphorylation. Phosphorylation of serines and tyrosines in Cxs have been shown to be essential for proper assembly and function of GJ channels (Lampe & Lau, 2004). To date, none of the other common PTMs have been studied in much details for Cxs. Studies aimed at identifying these different PTMs may be instrumental in understanding important aspects of channel trafficking and molecular mechanisms of channel regulation as well as GJ assembly. Some studies have highlighted the association of Cxs with lipids, which are most likely to have structural and/or functional roles. Channels composed of Cx32 have been found to be tightly associated with endogenous phospholipids, suggesting that such association might have Cx-specific regulatory and/or structural interactions with lipid membranes (Locke & Harris, 2009). Studies using mass spectrometric analysis of Cx32 in HeLa cells and rodent livers have identified new PTMs of Cx32. These modifications included hydroxylation and/or phosphorylation in the amino-terminal domain of Cx32, γ-carboxyglutamation in the cytoplasmic loop of both Cxs, phosphorylation and palmitoylation in its C-terminal tail (CT) (Locke et al., 2006). These PTMs may likely to be involved in regulation of Cx trafficking and assembly or functional modulation.

The present study was aimed at determining if Cx32 is modified by palmitoylation and the functional implications of this PTM in regulating the trafficking, assembly and/or stability of Cx32 was determined. Protein palmitoylation, a reversible post-translational lipid modification is an important mechanism for regulating protein subcellular localization, trafficking, translocation to lipid rafts, and stability (Blaskovic et al., 2013; Linder & Deschenes, 2007). Palmitoylation has no sequence requirement except a cysteine residue, normally found in the proximity of transmembrane helices. A sequence search revealed three cysteine residues in the CT of Cx32 which can be potentially palmitoylated. The findings from this study showed that Cx32 is palmitoylated at cysteine residue 217 in cell lines derived from prostate tumors (LNCaP cells). Surprisingly, this palmitoylation did not affect the trafficking of Cx32 or its ability to form GJs at the cell surface. However, the findings reveal the importance of two of these cysteine residues (C280, C283), potentially as part of a CAAX motif in the CT of Cx32, in regulating its trafficking from the Golgi and also its stability, thereby affecting its assembly into GJs.

## RESULTS

### Single cysteine mutants traffic to the cell surface normally and assemble into gap junctions

The CT of Cx32 harbors three cysteine residues. These cysteines are located at residues 217, 280 and 283 (**Figure 1A**). A great number of Cx32 mutations have been described associated with the genetic disease X-linked Charcot-Marie-Tooth disease (CMTX), affecting most portions of Cx32. Of these C217stop and C280G have also been reported (Bergoffen et al., 1993; Kleopa et al., 2012). To explore the role of these cysteines in regulating assembly of Cx32 into GJs, Cx32 mutants were constructed where the cysteine residues were mutated to alanines. These mutants are Cx32-C217A, where C217 is mutated to alanine, Cx32-C280A, where C280 is mutated to alanine, and Cx32-C283A, where C283 is mutated to alanine (**Figure 1B**). The fate of these mutants was explored in LNCaP cells, which do not express Cx32 (Katoch et al., 2015; Mehta et al., 1999). Cx32-WT, Cx32-C217A, Cx32-280A or Cx32-C283A was retrovirally expressed in LNCaP and pooled populations of these cells were obtained. Formation of GJs was then examined immunocytochemically. Cx32-WT expression was robust and the protein formed GJs, as seen in previous studies (Ray et al., 2018). β-catenin was used to delineate cell-cell interfaces. Results showed that similar to Cx32-WT, all mutants formed GJs (**Figure 2A**). Western blot analysis showed robust expression of Cx32-WT and the mutants (**Figure 2B**).

**Figure 1.**
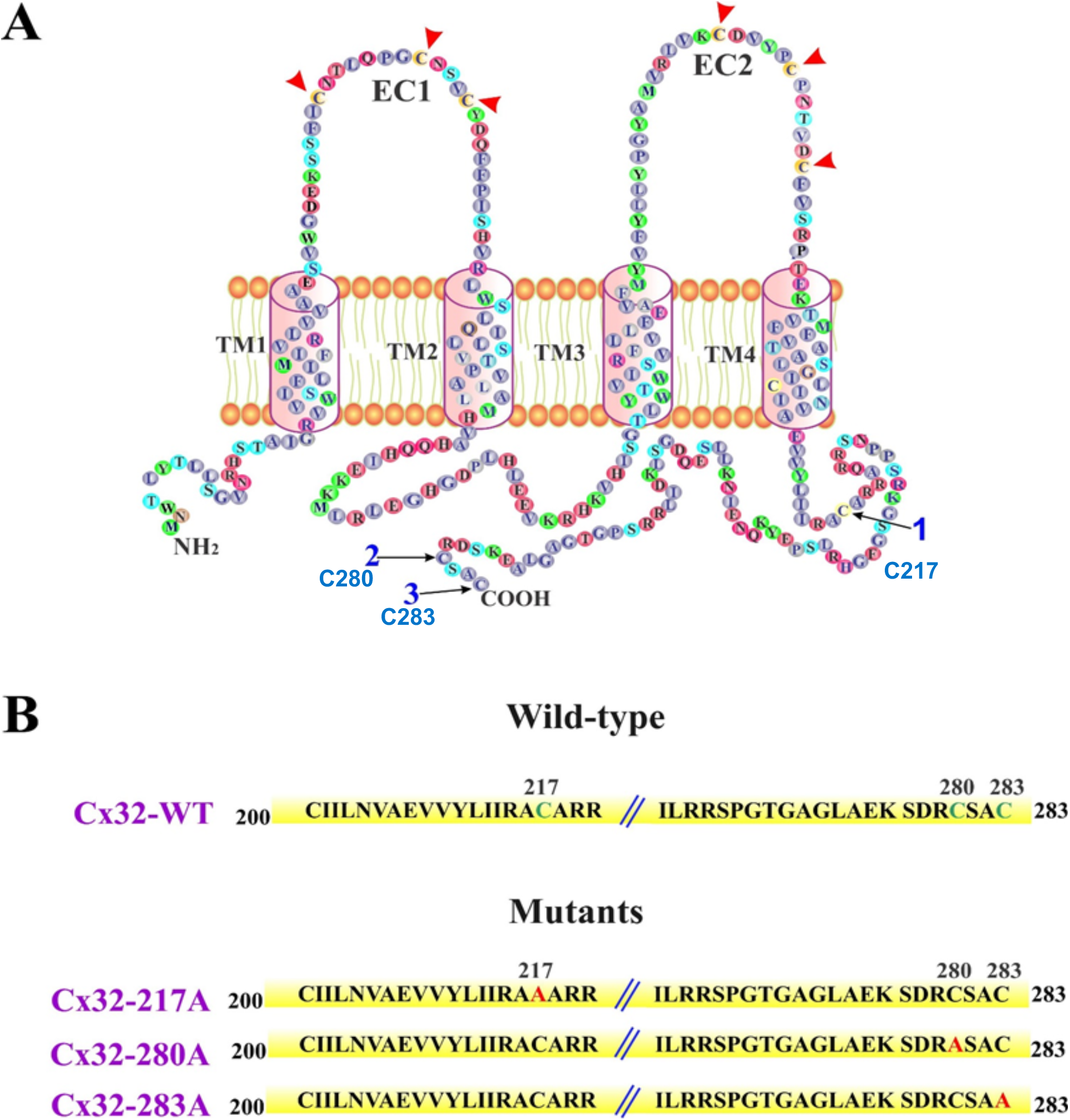
The single cysteine mutants of Cx32. **A**. Topology of Cx32. TM1-TM4 are the transmembrane domains. EC1 and EC2 are the extracellular loop domains. NH2 and COOH are the amino and the carboxyl termini, respectively. A single letter denotes an amino acid. The position of cysteine residues in the CT is indicated by the numbered arrows pointing towards them. Note also that there are six additional cysteines (3 in EC1 and 3 in EC2), which are not accessible to enzymatic modifications. They are indicated by the red arrowheads. **B**. Top: Position of cysteine residues that are likely to be lipidated in Cx32’s CT. The numbers on left and right indicate amino and carboxyl termini, respectively. The cysteine residues are shown in green. The numbers in black above the cysteine residues indicate their positions. Bottom: Mutants generated. The mutated amino acids are indicated in red. The names of mutants are shown in purple on the left. C=cysteine and A=alanine. In Cx32-C217A, cysteine at position 217 was mutated to alanine. In Cx32-C280A, cysteine at positions 280 was mutated to alanine and in mutant Cx32-C283A, cysteine at position 283 was mutated to alanine.

**Figure 2:**
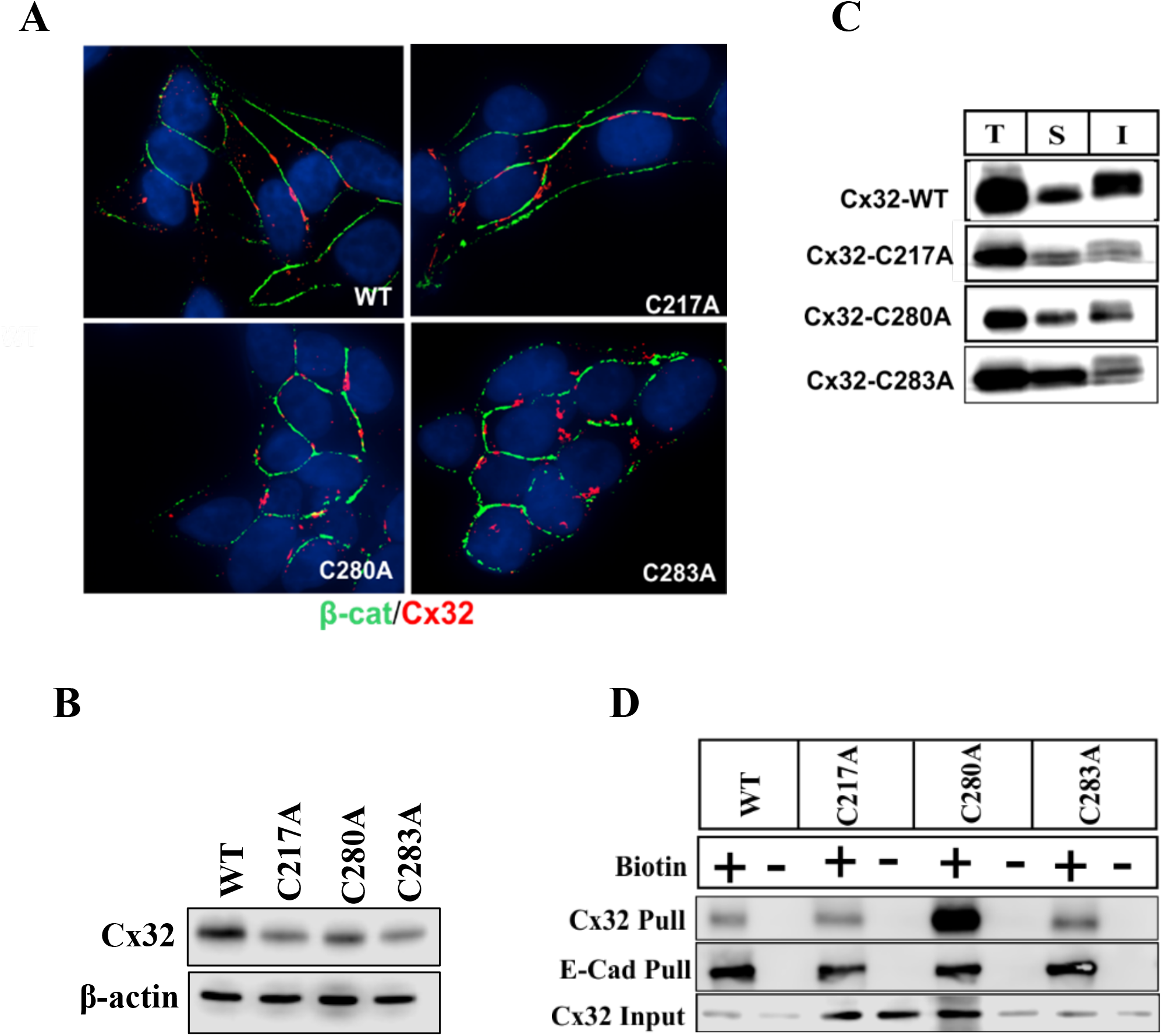
Single cysteine mutants form gap junctions and traffic to the cell surface normally. **A**. LNCaP cells expressing Cx32-WT or the mutants were immunostained for Cx32 (red) and β-catenin (β-cat, green). Note that the mutants efficiently assembled into GJs at cell-cell interfaces as pointed by white arrows. Nuclei are in blue. **B**. Western blot analysis of total cell lysates from cells expressing Cx32-WT or the mutants, showing robust and near equal expression. **C**. Detergent-solubility of Cx32-WT and the mutants. Western blot analysis of Cx32 in Total (T), TX100-soluble (S), and TX100-insoluble (I) fractions from LNCaP cells expressing Cx32-WT or the mutants, shows a predominantly insoluble fraction for all. **D**. Trafficking of Cx32-WT and single cysteine mutants. LNCaP cells expressing Cx32-WT, Cx32-C217A, Cx32-C280A or Cx32-C283A were cell-surface biotinylated (Biotin +). Biotinylated proteins were pulled down by streptavidin (Pull) and immunoblotted for Cx32 and E-cadherin (E-Cad). A non-biotinylated dish was kept as a control (Biotin -). Note that like Cx32-WT, all the mutants were efficiently biotinylated. For the input, 10 μg of total protein was used and immunoblotted for Cx32.

To corroborate the immunocytochemical data, the GJ forming ability of Cx32-WT, Cx32-C217A, Cx32-280A and Cx32-C283A was examined biochemically by the detergent-solubility assay using 1% Triton X-100 (TX100). Analyses of total as well as TX100-soluble and -insoluble fractions from cells expressing Cx32-WT or the mutants by Western blotting showed that GJs composed of Cx32-WT, Cx32-C217A, Cx32-280A or Cx32-C283A appeared predominantly in the detergent-insoluble fraction (**Figure 2C**), suggesting that Cx32-WT and the mutants assembled into GJs.

To assess whether Cx32-WT and the mutants trafficked normally to the cell surface in LNCaP cells, a cell surface biotinylation assay was performed. The results showed that Cx32-WT and the mutants were robustly biotinylated. There was no difference in the biotinylation of E-cadherin, a cell surface protein, which was used as a positive control (**Figure 2D**). Taken together, these results suggested that mutating C217, C280 or C283 singly to alanines, affected neither the trafficking of Cx32 to the cell surface nor its ability to assemble into GJs.

### The triple cysteine mutant of Cx32 fails to assemble into gap junctions

Because single cysteine mutants of Cx32 assembled into GJs (**Figure 2**) and multiple cysteines could be lipidated in a given protein, the effect of mutating all three cysteines on GJ assembly was next examined. Hence, a mutant Cx32 was constructed where C217, C280 and C283 together were mutated to alanines. This triple mutant was referred to as Cx32-3CA (**Figure 3A**). Cx32-WT or Cx32-3CA were retrovirally expressed in LNCaP cells. Pooled polyclonal cultures were next analyzed to assess GJ assembly. Surprisingly, immunocytochemical analysis revealed that as compared to Cx32-WT, which formed large GJs, Cx32-3CA did not and instead accumulated intracellularly (**Figure 3B**). Moreover, Western blot analysis showed that the expression level of Cx32-3CA was lower than that of Cx32-WT (**Figure 3C**). To substantiate the immunocytochemical data, we examined the GJ forming ability of Cx32-WT and Cx32-3CA biochemically by the detergent-solubility assay using 1% TX100. Analyses of total as well as TX100-soluble and -insoluble fractions from cells expressing Cx32-WT or Cx32-3CA by Western blotting showed that while GJs composed of Cx32-WT were found in the detergent-insoluble fraction, those formed of Cx32-3CA were found predominantly in the soluble fraction (**Figure 3D**). This suggested that Cx32-3CA fails to assemble into GJs and the intracellular puncta were not detergent-insoluble denatured aggregates of Cx32-3CA.

**Figure 3:**
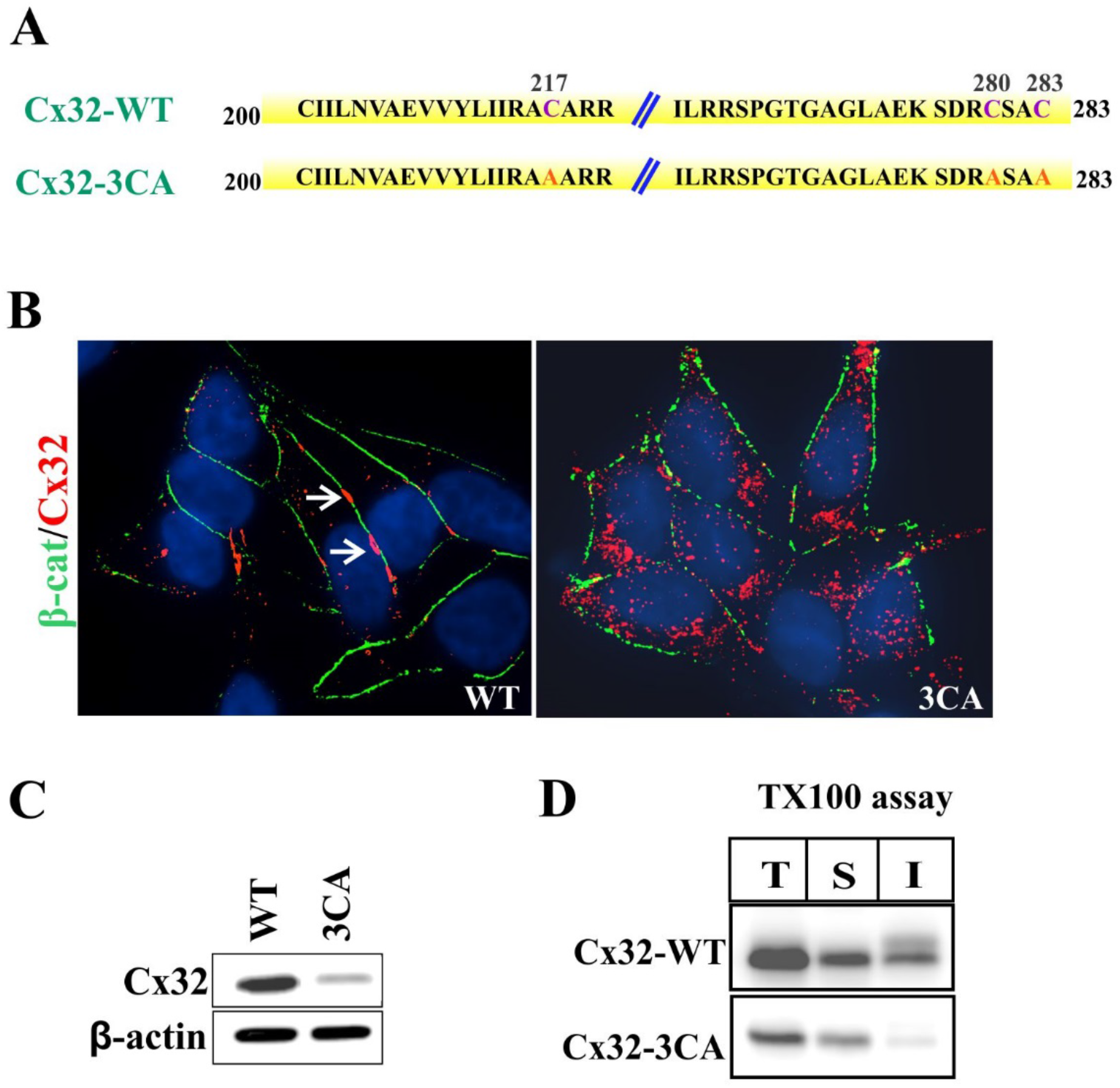
The triple cysteine mutant of Cx32 (Cx32-3CA) accumulates intracellularly and does not assemble into gap junctions. **A**. The position of the cysteine residues in the CT of Cx32-WT. The numbers on left and right indicate amino and carboxyl termini, respectively. The cysteine residues (C) are shown in purple in the CTs of Cx32-WT. The numbers in black above the cysteine residues indicate the position of the amino acids. Bottom: Mutant Cx32-3CA. The mutated amino acids (mutated to alanine, A) are indicated in red. **B**. LNCaP cells expressing Cx32-WT or Cx32-3CA were immunostained for Cx32 (red) and β-catenin (β-cat, green). Note that Cx32-3CA did not assemble into GJs whereas Cx32-WT did (indicated by white arrows) and was localized intracellularly. Nuclei are in blue. **C**. Western blot analysis of total cell lysates from cells expressing Cx32-WT or Cx32-3CA. **D**. Detergent-solubility of Cx32-WT and Cx32-3CA. Western blot analysis of Cx32 in total (T), TX100-soluble (S), and TX100-insoluble (I) fractions from LNCaP cells expressing Cx32-WT or Cx32-3CA. Note that Cx32-3CA was found predominantly in the detergent-soluble fraction whereas Cx32-WT was predominantly found in the detergent-insoluble fraction.

### Assembly of Cx32 into gap junction is independent of gap junction assembly

Palmitoylation plays an important role in influencing protein trafficking, protein stability, and protein subcellular localization (Linder & Deschenes, 2007). Earlier studies using mass spectrometric analysis using mouse Cx32 had detected a palmitoylated C-terminal peptide (residues 277–283) where either C280 or C283 or both could be palmitoylated (Locke et al., 2006). A database search using SwissPalm was performed to assess the palmitoylation state of Cx32. SwissPalm is a protein palmitoylation database which integrates predictions of S-palmitoylated cysteine scores, orthologs and isoform multiple alignments to predict the palmitoylation status of a protein (Blanc et al., 2015). Analysis using SwisssPalm showed a high prediction confidence for C217 and C280 to be palmitoylated (**Figure 4A**). Next was examined whether the palmitoylation of Cx32 regulates its trafficking and assembly into GJs. To test this, it was first assessed if Cx32 was palmitoylated and which cysteines were involved. To investigate this, the Acyl Biotin Exchange (ABE) pulldown assay, was performed which is a well-established method to detect protein palmitoylation (Wan et al., 2007) (See Materials and Methods). In this assay, Cx32-3CA was used as a negative control, which would not be palmitoylated as all the cysteines in the CT are mutated to alanines. Results showed that Cx32-WT, Cx32-C280A and Cx32-C283A were palmitoylated whereas Cx32-C217A and Cx32-3CA were not (**Figure 4B**), thereby suggesting that Cx32 is palmitoylated at C217. To assess if palmitoylation is regulating the trafficking of Cx32, we treated LNCaP cells expressing Cx32-WT with 100 μM 2-Bromopalmitate (2-BP), an inhibitor of palmitoylation (Webb et al., 2000). Cells were co-immunostained with Golgi-marker, TGN46, to check if blocking palmitoylation disrupts trafficking of Cx32-WT to the cell surface and cause its trafficking arrest in the Golgi. Results showed that in 2-BP treated cells, Cx32-WT was able to traffic and form GJs normally, and no enhanced colocalization was observed with TGN46 (**Figure 4C**). To further assess if defect in palmitoylation affects GJ size, the surface areas of GJs at several cell-cell interfaces was measured (see Materials and Methods). Results showed no difference between Cx32-WT and Cx32-C217A (**Figure 4D**) in terms of surface area. Because C217A formed GJs and trafficked normally (**Figure 2**), and blocking palmitoylation did not affect trafficking or assembly of Cx32-WT (**Figure 4C, D**); these data implied that palmitoylation does not play a role in regulating the trafficking of Cx32 and/or its assembly into GJs.

**Figure 4:**
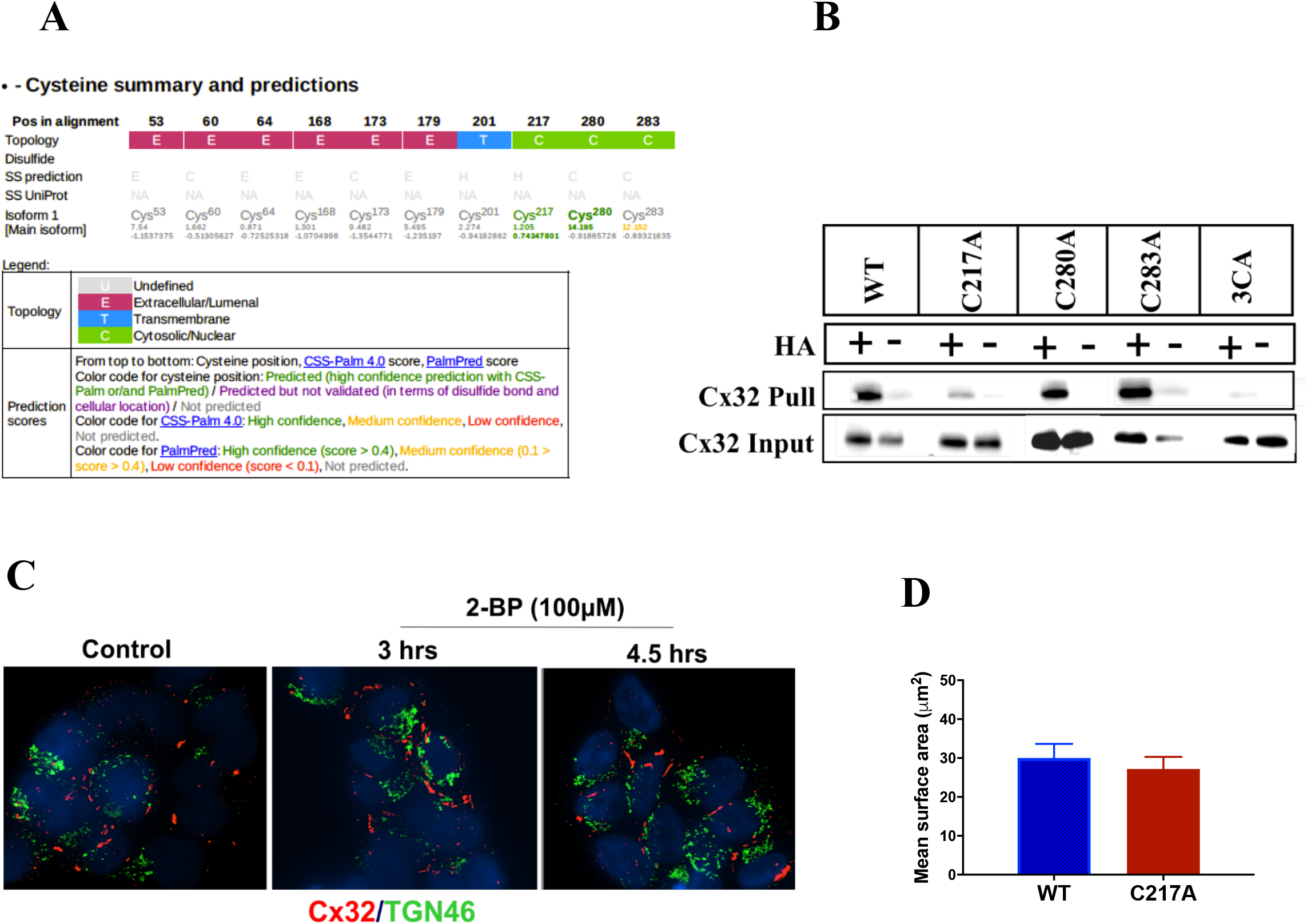
Assembly of Cx32 into gap junctions occur independently of palmitoylation. **A**. A search using the SwissPalm database indicated a high confidence prediction score for palmitoylation at C217 and C280, both of which are cytosolic and present in the CT of Cx32. Code for cysteine position E=Extracellular, T=Transmembrane, C-Cytosolic. Color code for prediction: High confidence, Medium confidence, Low confidence, Not predicted. Note, C217 and C280 show a high confidence prediction. **B**. ABE Pulldown assay. Lysates from LNCaP cells expressing Cx32-WT, Cx32-C217A, Cx32-C280A, Cx32-C283A or Cx32-3CA were split into two halves and treated with HA (+) or left untreated (-). After the streptavidin-agarose pull-down, the captured proteins were resolved by SDS-PAGE and immunoblotted for Cx32. (See Materials and Methods). For the input, 10 μg of total protein was used and immunoblotted for Cx32. Note that only the C217A mutant was palmitoylated. **C**. LNCaP cells expressing Cx32-WT were subjected to 100 μM 2-BP treatment for 3h and 4.5h. Cells were then fixed and immunostained for Cx32 (red) and TGN46 (green). **D**. GJ areas of cells expressing Cx32-WT or the mutant C217. Surface areas (Mean ± SEM) of 40-70 distinct GJ puncta at cell-cell interfaces from 2 independent experiments were determined using the measurement module of Volocity. The area is represented in μm^2^. Note no change between the mean GJ area in cells expressing the mutant C217A and the WT.

### Intracellular accumulation is due to double mutation at C280 and C283

Previous results showed that the single cysteine Cx32 mutants trafficked to the cell surface and assembled into GJs (**Figure 2**), whereas Cx32-3CA, where all cysteines were mutated, accumulated intracellularly (**Figure 3**). Therefore, it was further explored which of the three cysteines, in combination of two, were responsible for intracellular accumulation. To assess this, a series of double cysteine mutants of Cx32 were constructed as follows: Cx32-C217A/C280A in which cysteine residues at positions 217 and 280 were mutated to alanines, Cx32-C217/C283A in which cysteine residues at positions 217 and 283 were mutated to alanines, and Cx32-C280A/C283A in which cysteine residues at positions 280 and 283 were mutated to alanines (**Figure 5A**). These double mutants were then retrovirally expressed in LNCaP cells in parallel. Their ability to form GJs was assessed immunocytochemically whereas their expression was assessed by Western blotting. Immunocytochemical analysis showed that both Cx32-C217A/C280A and Cx32-C217/C283A formed GJs at cell-cell interfaces, whereas Cx32-C280A/C283A did not, and accumulated intracellularly (**Figure 5B, top panels**), similar to Cx32-3CA (**Figure 3B**). To substantiate the immunocytochemical data, these mutants were expressed in another cell line HEK293T. Cx32-WT or the double cysteine mutants were transiently transfected into HEK293T and immunostained for Cx32 after 24 h. As was observed in LNCaP cells, the mutant Cx32-C280A/C283A accumulated intracellularly whereas the other mutants were assembled into GJs like Cx32-WT (**Figure 5B, bottom panels**). Western blot analysis showed a reduced expression level of Cx32-C280A/C283A compared to Cx32-WT in LNCaP cells (**Figure 5C**), similar to that observed for Cx32-3CA (**Figure 3C**). Henceforth, the mutant Cx32-C280A/C283A will be referred to as Cx32-2CA. To further corroborate the immunocytochemical data, the GJ forming ability of Cx32-WT and Cx32-2CA was examined biochemically by the detergent-solubility assay using 1% TX100. Analyses of total as well as TX100-soluble and -insoluble fractions from cells expressing Cx32-WT or Cx32-2CA by Western blotting showed that Cx32-WT was found predominantly in the detergent-insoluble fraction, whereas Cx32-2CA occurred predominantly in the soluble fraction (**Figure 5D**). Thus, the detergent solubility assay confirmed that Cx32-2CA fails to assemble into GJs and that Cx32-2CA is not aggregated into detergent-insoluble puncta. Altogether, the above results suggested that intracellular accumulation is due to double mutation at C280 and C283.

**Figure 5:**
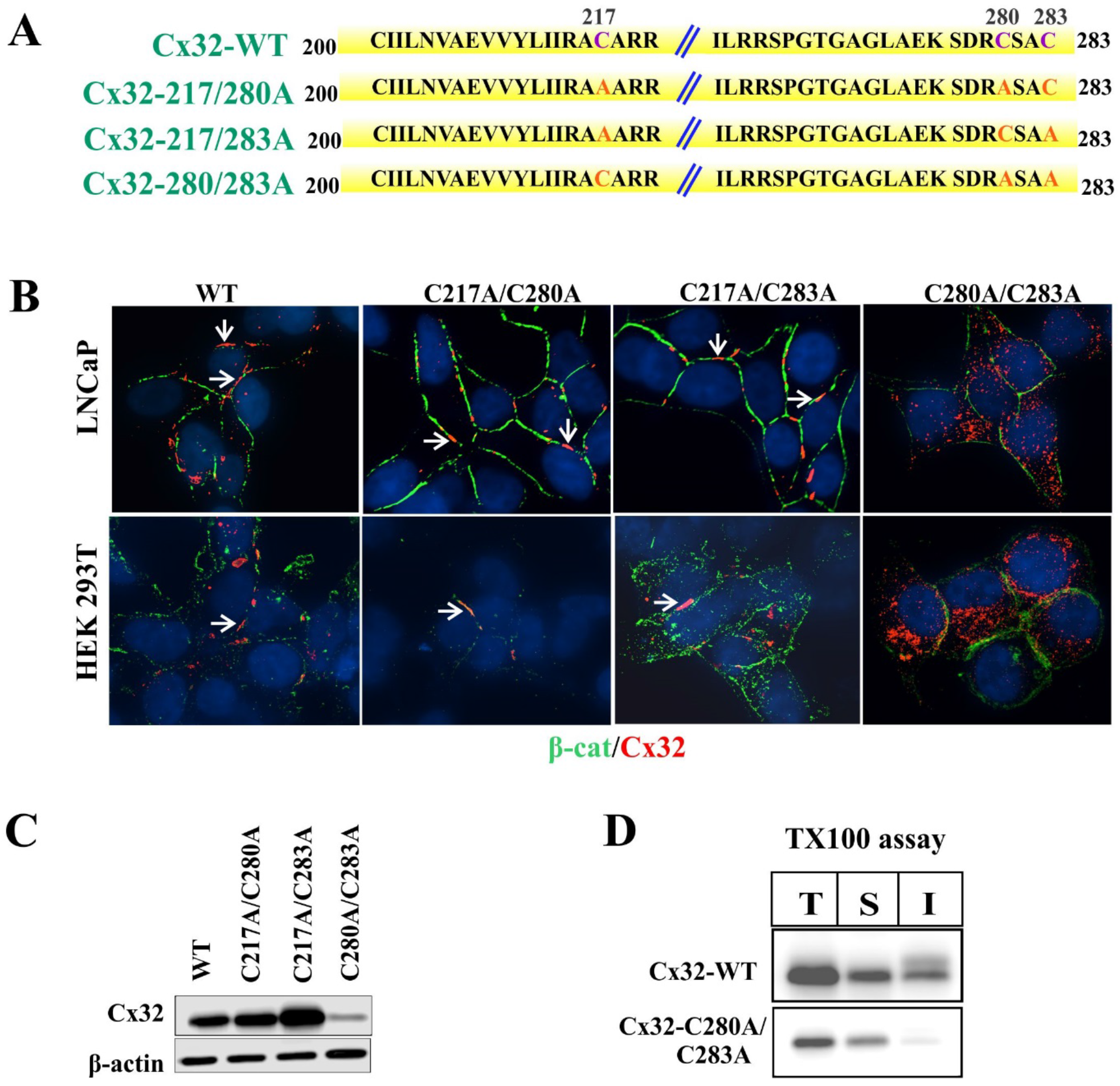
Intracellular accumulation is due to double mutation at C280 and C283. **A**. Different double cysteine mutant constructs of Cx32. The numbers on left and right indicate amino and carboxyl termini, respectively. The cysteine (C) residues are shown in purple in the CT of Cx32-WT. The numbers in black above the cysteine residues indicate the position of the amino acids. The names of the mutants generated are shown in green on the left. The mutated amino acids (mutated to alanine, A) are indicated in red. **B**. LNCaP cells (top panels) and HEK293T cells (bottom panels) expressing Cx32-WT and the mutants were immunostained for Cx32 (red) and β-catenin (β-cat, green). Note that Cx32-WT, Cx32-C217A/C280A and Cx32-C217A/C283A assembled into GJs (indicated by white arrows), whereas Cx32-C280A/C283A did not, and was localized intracellularly in both LNCaP and HEK293T cells. Nuclei are shown in blue. **C**. Western blot analysis of total cell lysates from cells expressing Cx32-WT and Cx32-C280A/C283A. Note reduced expression level of Cx32-C280A/C283A compared to Cx32-WT and other mutants. **D**. Detergent-solubility of Cx32-WT and Cx32-C280A/C283A. Western blot analysis of Cx32 in Total (T), TX100-soluble (S), and TX100-insoluble (I) fractions from LNCaP cells expressing Cx32-WT or Cx32-3CA. Note a significant detergent-soluble fraction in cells expressing Cx32-3CA.

To assess if lack of palmitoylaton in Cx32-2CA is the cause of defective trafficking, an ABE assay was performed with LNCaP cells expressing Cx32-2CA, using Cx32-WT expressing LNCaP cells as our positive control and Cx32-3CA expressing LNCaP cells as our negative control. In Cx32-2CA, only C217, which was found to be palmitoylated (**Figure 4B**), is intact. If mutating C280A and C283A together is abolishing palmitoylation on C217, then this impaired trafficking of Cx32-2CA can be attributed to a lack of palmitoylation. Results showed that similar to Cx32-WT, Cx32-2CA was also palmitoylated, whereas Cx32-3CA was not (**Figure S1**). This data suggested that palmitoylation in Cx32-2CA is not affected by mutating the two terminal cysteines C280A and C283A, thereby ruling out lack of palmitoylation as a cause of impaired trafficking.

Together, the above results are suggestive that Cx32-2CA and Cx32-3CA behave similarly with respect to their protein expression, intracellular accumulation, impaired trafficking and inability to assemble into GJs, and the above processes are independent of palmitoylation and the double mutation at C280 and C283 being the cause. Hence, in our subsequent experiments we have used Cx32-3CA to serve as an additional control for the results obtained for Cx32-2CA.

### Intracellular accumulation of Cx32-2CA and Cx32-3CA is due to impaired trafficking to the cell surface from the Golgi

The intracellular accumulation of Cx32-2CA and Cx32-3CA could be due to two possibilities: their defective trafficking from the trans-Golgi network (TGN) to the cell surface or rapid endocytosis from the cell surface prior to assembly. To test the first possibility, a cell-surface biotinylation assay was performed to determine whether impaired trafficking of Cx32-2CA and Cx32-3CA compared to Cx32-WT might account for their failure to form GJs. Results showed that compared to Cx32-WT, Cx32-2CA and Cx32-3CA were poorly biotinylated (**Figure 6A**). There was no difference in the biotinylation of E-cadherin, a cell surface protein, which was used as a positive control. Therefore, this data suggested that both the mutants trafficked to the cell surface poorly.

**Figure 6:**
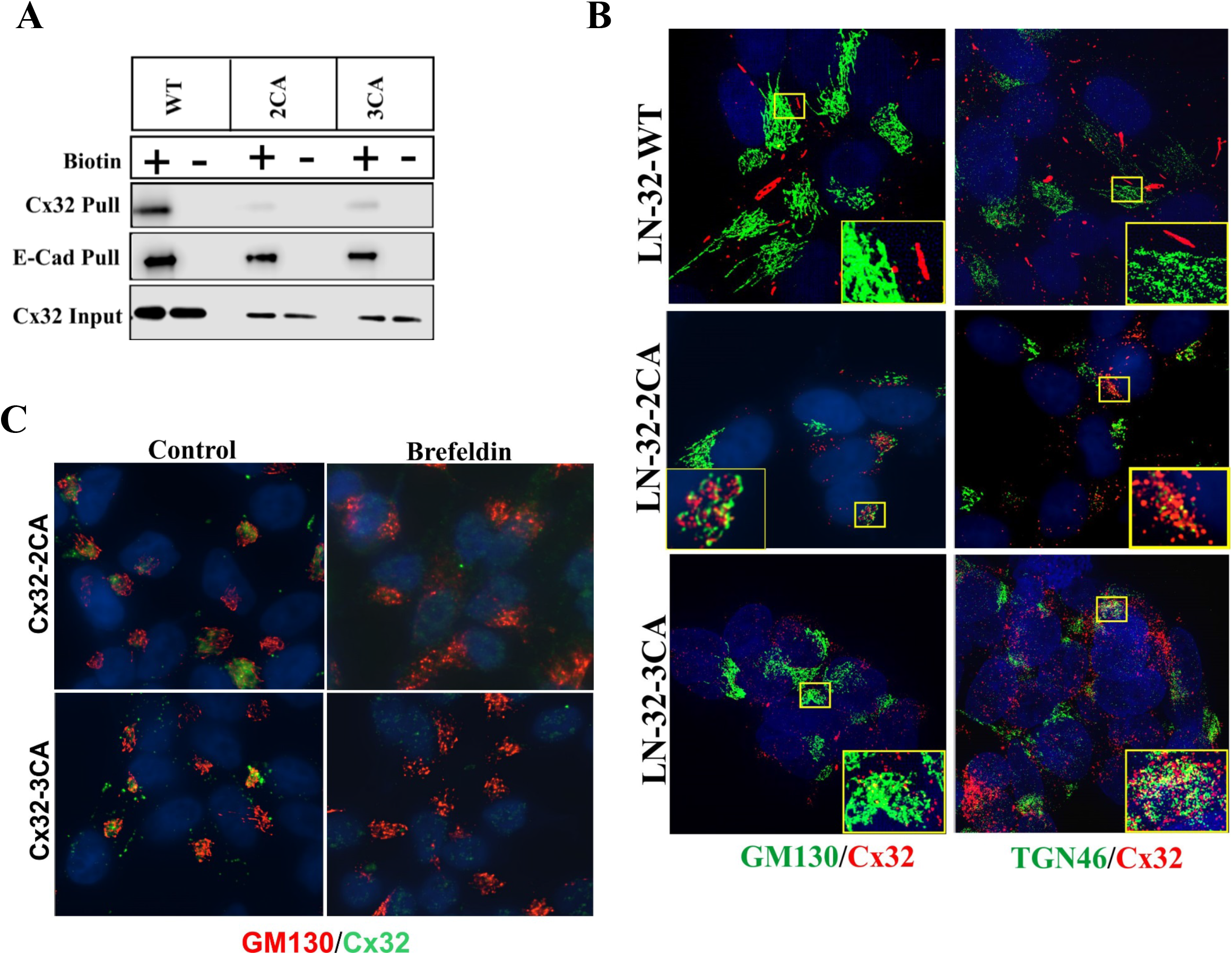
Intracellular localization of Cx32-2CA and Cx32-3CA is due to impaired trafficking to the cell surface. **A**. LNCaP cells expressing Cx32-WT, Cx32-2CA or Cx32-3CA were cell-surface biotinylated (Biotin +). Biotinylated proteins were pulled down by streptavidin (Pull) and immunoblotted for Cx32 and E-cadherin (E-Cad). A non-biotinylated dish was kept as a control (Biotin -). Note that compared to Cx32-WT, mutant Cx32-2CA and Cx32-3CA were poorly biotinylated. Note also that E-cadherin was equally biotinylated in cells expressing Cx32-WT, Cx32-2CA or Cx32-3CA. Note also reduced expression level of Cx32-2CA and Cx32-3CA compared to Cx32-WT. For the input, 10 μg of total protein was used and immunoblotted for Cx32. **B**. LNCaP cells expressing Cx32-WT, Cx32-2CA or Cx32-3CA were immunostained for Cx32 (red) and GM130 or TGN46 (green). The enlarged images of boxed areas in the panels are shown at the bottom right. Note robust colocalization of the mutants with the indicated markers as compared to Cx32-WT. **C**. LNCaP cells expressing Cx32-2CA or Cx32-3CA were treated with vehicle (Control) or 10 μM brefeldin A (Brefeldin) for 8 h. Cells were then immunostained for Cx32 (green) and GM130 (red). Note loss of intracellular localization pattern of Cx32-2CA and Cx32-3CA upon brefeldin treatment.

Since Cx32-2CA and Cx32-3CA showed predominantly an intracellular pattern of localization (**Figure 3B, 4B**), LNCaP cells expressing Cx32-2CA or Cx32-3CA were next immunostained with Golgi markers, GM130, a cis-Golgi marker (Nakamura et al., 1995); and TGN46, a marker for the TGN (De Matteis & Luini, 2008), to assess the localization of these mutants in the cell. Robust colocalization of Cx32-2CA and Cx32-3CA with the above Golgi markers was observed as compared to Cx32-WT (**Figure 6B**), suggesting that these mutants reside in the Golgi or traffick through the Golgi *enroute* to the cell surface. To support these observations and to rule out that this intracellular localization of Cx32-2CA or Cx32-3CA was unique to LNCaP cells only, this phenotype was confirmed in different cell lines. Cx32-WT, Cx32-2CA or Cx32-3CA was retrovirally expressed in BxPC3 cells (**Figure S2A**); and transiently expressed in HEK293T cells (**Figure S2B**). Their localization was then assessed immunocytochemically by co-immunostaining with cell-surface marker, β-catenin, and Golgi marker, TGN46. Results showed that this intracellular pattern of localization of Cx32-2CA and Cx32-3CA was also maintained in BxPC3 and HEK293T cells and these mutants failed to form GJs (**Figure S2**). To further confirm the above results, the localization pattern of Cx32-2CA and Cx32-3CA was examined upon treatment with the antiviral antibiotic brefeldin A, which is known to cause disassembly of the Golgi complex and redistribution of Golgi proteins to the ER, thereby causing a trafficking defect from the ER to the Golgi (Fujiwara et al., 1988). For these experiments, LNCaP cells expressing these mutants were treated with brefeldin (10 μM) for 8 h and it was observed that the intracellular pattern of localization was almost lost upon treatment (**Figure 6C**). Altogether, these results suggested that these mutants predominantly reside in the Golgi and their defective phenotype is due to a trafficking defect from the Golgi to the cell surface

Next was examined whether this lack of cell-surface biotinylation and intracellular accumulation of Cx32-2CA and Cx32-3CA was caused due to rapid endocytosis of these mutants followed by retrograde transport back to the Golgi. If so, inhibiting endocytosis would induce the assembly of these mutants into GJs. To test this notion, cells were subjected to potassium (K^+^) depletion or hypertonic sucrose (Suc). Both treatments inhibit clathrin-mediated endocytosis (Hansen et al., 1993; Larkin et al., 1986). Alexa 594-conjugated transferrin receptor (Tfr) was used as a positive control for clathrin-mediated endocytosis. As expected, Alexa 594-conjugated Tfr was internalized in control cells but not in cells subjected to K^+^ depletion or hypertonic sucrose (**Figure 7AB, top panels**). However, potassium (K^+^) depletion or treating cells with hypertonic sucrose for 2 h did not induce the assembly of Cx32-2CA or Cx32-3CA into GJs as assessed immunocytochemically (**Figure 7AB, bottom panels**). Taken together, these results suggested that the Golgi accumulation of Cx32-2CA and Cx32-3CA was caused by impaired trafficking from the Golgi to the cell surface, and not by their endocytosis prior to assembly into GJs.

**Figure 7.**
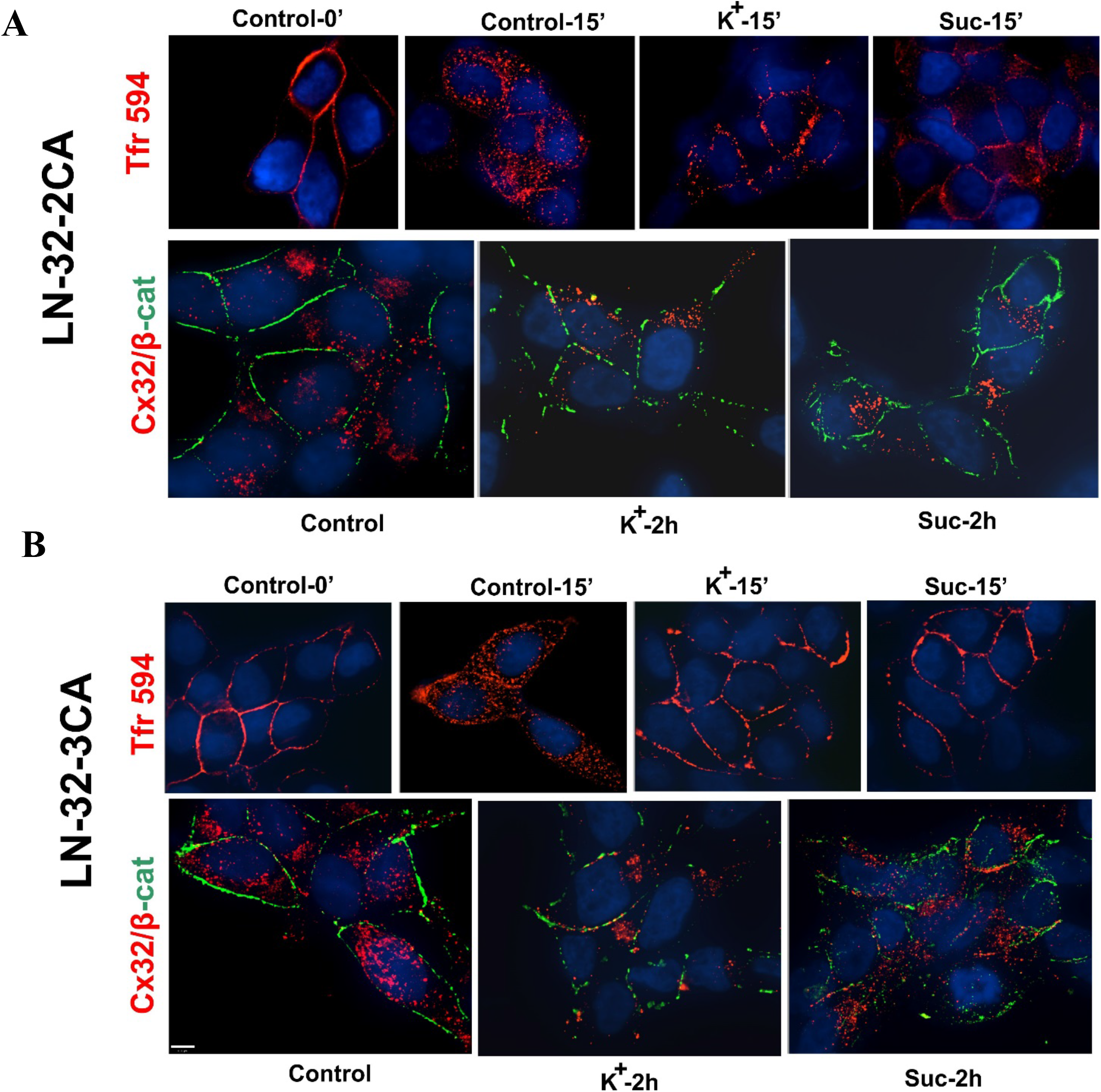
32-2CA and Cx32-3CA mutants does not undergo endocytosis prior to assembly. LNCaP cells expressing (**A**) Cx32-2CA or (**B**) Cx32-3CA were subjected to K^+^ depletion (K^+^) or hypertonic sucrose (Suc) treatment for 2 h. Cells were then fixed and immunostained for Cx32 (red). As a control, these cells were labeled with Alexa Fluor 594-conjugated transferrin (Tfr) and were left untreated (Control) for 0 or 15 minutes or subjected to K^+^ depletion or Suc treatment for 0 or 15 minutes (0’ or 15’). Note internalization of Alexa Fluor 594-conjugated Tfr in control cells but not in cells subjected to K^+^ depletion or Suc treatment. Note also that the intracellular localization of Cx32-2CA or Cx32-3CA is not rescued upon K^+^ depletion or Suc treatment, suggesting that they do not undergo endocytosis prior to assembly.

### Cx32 cysteine mutants undergo rapid turnover due to enhanced lysosomal degradation

Assessing protein expression levels between Cx32-WT and Cx32-2CA or Cx32-3CA showed a 70-80% decrease in Cx32 protein levels in these mutants as compared to the wild-type, as shown by Western blot analysis (**Figure 8A**) and its quantification (**Figure 8B**). To assess if this significant decrease in protein expression levels was a result due to changes in the gene transcription levels, mRNA expression levels of Cx32 in the mutants compared to Cx32-WT were analyzed. Results showed a near equal mRNA expression level between Cx32-WT and Cx32-2CA or Cx32-3CA (**Figure 8C**). This suggested that the decrease of Cx32 protein levels observed in the Cx32-cysteine mutants (**Figure 8A**) was not the result of decreased mRNA levels but possibly due to an increased degradation at the protein level as compared to Cx32-WT. The half-life of Cxs is short and ranges between 2-5 h *in vivo* and *in vitro* (Beardslee et al., 1998; Fallon & Goodenough, 1981; VanSlyke et al., 2000). To test the stability of these mutants, the wild-type and mutant Cx32 expressing LNCaP cells were treated with cycloheximide (CHX), a protein synthesis inhibitor and CHX chase assay was performed. LNCaP cells expressing Cx32-WT, Cx32-2CA or Cx32-3CA were grown in media supplemented with 60 μM CHX over a period of several hours, lysates were collected at different time-points and immunoblotted with anti-Cx32 antibody. Within 4 hours, Cx32-2CA and Cx32-3CA showed almost 100% degradation, whereas Cx32-WT levels reduced by approximately 50% only (**Figure 8D**). Cx32-2CA and Cx32-3CA degraded with a half-life (t_1/2_) of approximately one hour, in contrast to Cx32-WT which had a t_1/2_ of approximately 5 hours. Immunocytochemical analysis showed similar results, where Cx32 staining (shown in red) decreased considerably around 2 h for Cx2-2CA and Cx32-3CA, with almost no signal observed at 4 h, in contrast to Cx32-WT, upon treatment with CHX (**Figure 8E**). Hence, these results suggested that the reduced proteins levels of Cx32-2CA and Cx32-3CA were a result of enhanced protein degradation.

**Figure 8.**
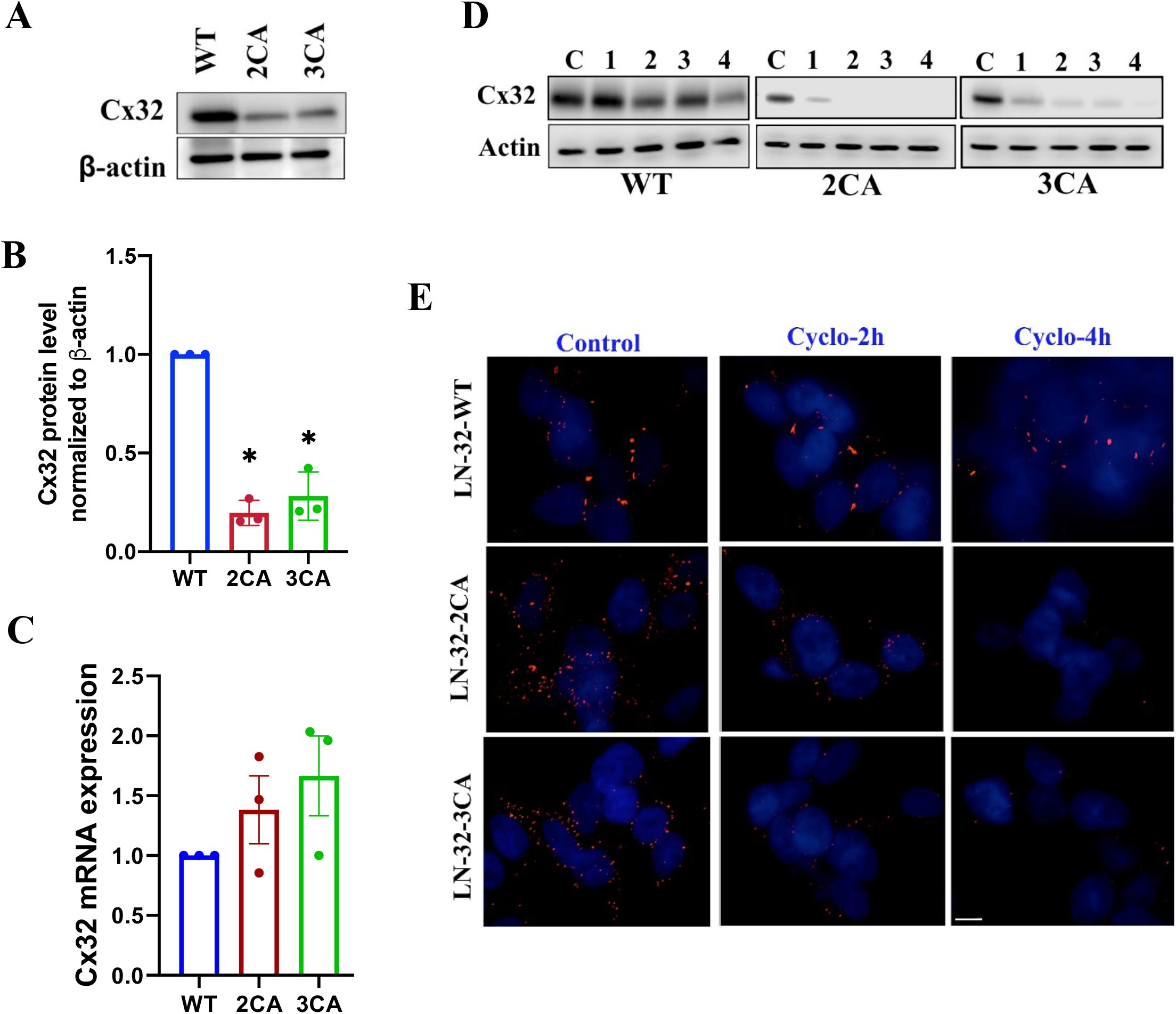
Cx32 cysteine mutants are downregulated post-translationally. **A**. Western blot analysis of total cell lysates from LNCaP cells expressing Cx32-WT, Cx32-2CA or Cx32-3CA. β-actin is used as a loading control. **B**. Quantification of the Western Blot data from 3 independent experiments, represented as Cx32 levels relative to β-actin levels. n=3 ± SEM, normalized to the WT. * p< 0.05. Each data point represents an individual experiment. Note a significant reduction in protein expression levels for Cx32-2CA or Cx32-3CA, when compared to Cx32-WT. **C**. Quantification of mRNA expression levels of LNCaP ells expressing Cx32-WT, Cx32-2CA or Cx32-3CA, relative to 36B4 (internal control) by RT-PCR, n=3 ± SEM, normalized to the WT. Each data point represents an individual experiment. Note no significant change in the mRNA expression levels for Cx32-2CA or Cx32-3CA, when compared to Cx32-WT. **D**. Western blot analysis of lysates from confluent LNCaP cells expressing Cx32-WT, Cx32-2CA or Cx32-3CA subjected to a CHX-chase assay for upto 4 hours using 60 μM CHX. C denotes treatment with vehicle (control). **E**. Immunocytochemical analysis of LNCaP cells expressing Cx32-WT, Cx32-2CA or Cx32-3CA subjected to a similar CHX-chase assay as described above and immunostained for Cx32 (red). Only 2h and 4h timepoints are shown. Note the decrease in Cx32 immunostaining at higher timepoints for Cx32-2CA and Cx32-3CA in contrast to Cx32-WT. 33

Next it was determined whether the reduction in protein levels were a result of proteosomal or lysosomal degradation. To test this, LNCaP cells expressing Cx32-WT, Cx32-2CA or Cx32-3CA were treated with leupeptin and ammonium chloride (lysosomal inhibitors) and lactacystin (proteosomal inhibitor). Western Blot analysis showed that treatment with lysosomal inhibitors enhanced the protein levels of Cx32-2CA and Cx32-3CA greatly, more than the increase observed with Cx32-WT. Note, that the levels of the cytoskeletal protein β-actin used as a control was not affected. On the other hand, proteosomal inhibition failed to show any increase in the protein level of Cx32-WT or the mutants (**Figure 9A)**. We corroborated the above results immunocytochemically. A marked increase in intracellular Cx32 staining of these mutants was observed upon treatment with the lysosomal inhibitors (**Figure 9B**). Interestingly, lactacystin treatment showed no increase in intracellular staining, but a marked increase in gap junction assembly in the cells expressing Cx32-2CA or Cx32-3CA. As lysosomal inhibition increased the protein levels of the mutants, the localization of these mutants in the lysosomes was next examined by co-immunostaining these cells with CathepsinD, a lysosomal marker, along with Cx32-WT, Cx32-2CA or Cx32-3CA under steady-state conditions as well as upon treatment with the above lysosomal inhibitors. Results showed discernible colocalization of the cysteine mutants with CathepsinD upon treatment with these inhibitors (**Figure 9C**). These results suggested that mutating the cysteines results in decreased stability of Cx32 by way of enhanced lysosomal degradation, and inhibiting lysosomal degradation rescues this reduced protein levels observed for these mutants.

**Figure 9.**
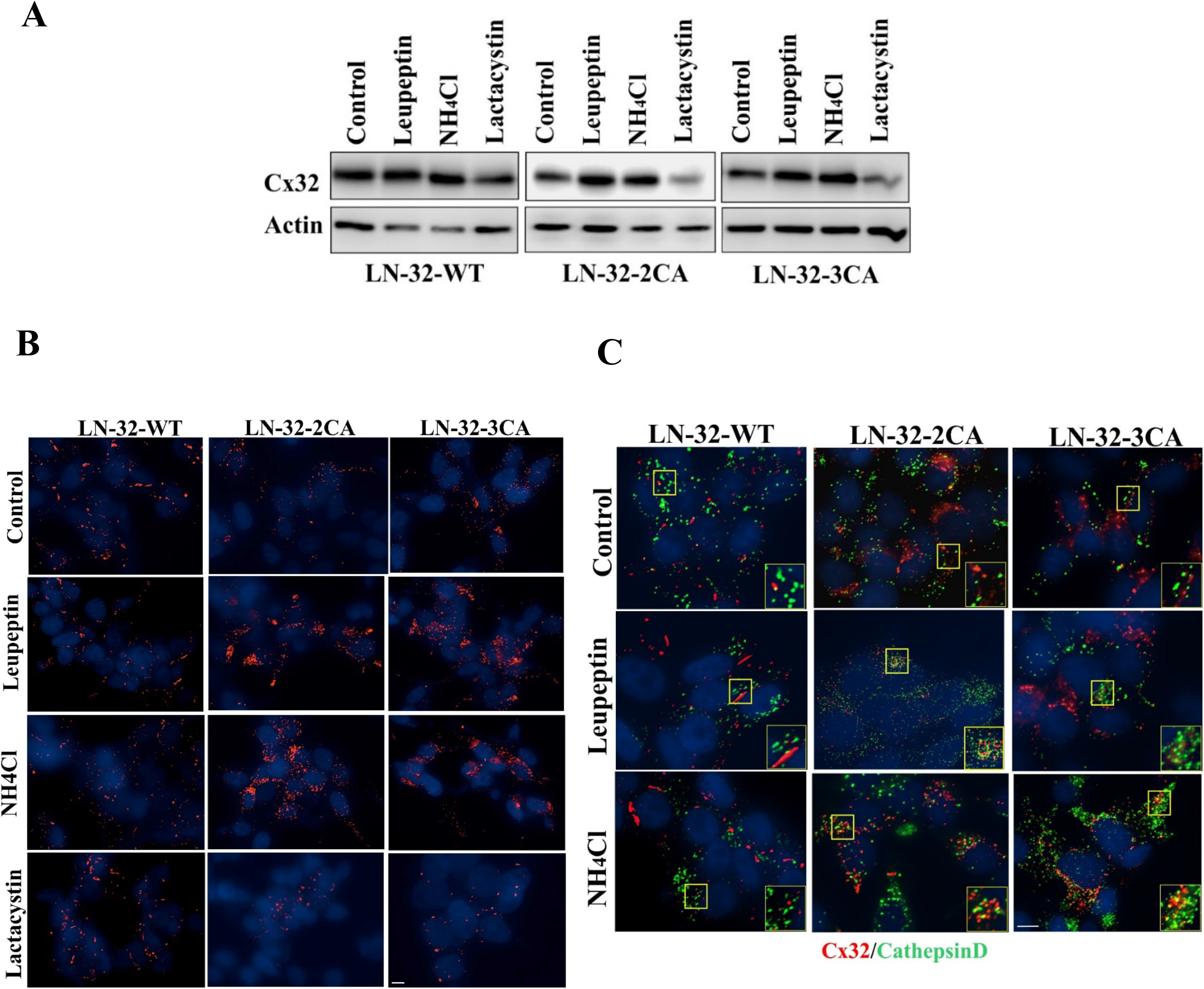
Cx32 cysteine mutants undergo rapid lysosomal degradation. **A**. LNCaP cells expressing Cx32-WT, Cx32-2CA or Cx32-3CA were either untreated (Control) or treated with 200 μM leupeptin for 8 h, 10 mM NH_4_Cl for 6 h and 10 μM lactacystin for 6h. Expression level of Cx32 and mutants was determined by **A**. Western blot analysis. **B**. Immunocytochemically by staining for Cx32 (in red). Note inhibition of degradation of Cx32-WT and the mutants upon leupeptin and NH_4_Cl treatment. **C**. The above cells were coimmunostained with CathepsinD (green) and Cx32 (red). Note discernible colocalization of Cx32-2CA and Cx32-3CA with CathepsinD upon treatment with leupeptin and NH_4_Cl. Scale bar = 10 μm.

## DISCUSSION

Connexins are subjected to a wide variety of PTMs. However, not much has been studied about how different PTMs, except phosphorylation, can regulate Cx trafficking and assembly. Most of these PTMs occur in the CT of Cxs. Previous studies showed that Cx32-CT is palmitoylated (Huang et al., 2005; Locke et al., 2006) and this PTM plays a significant role in regulating the trafficking, stability and cellular localization for many other proteins (Blaskovic et al., 2013; Linder & Deschenes, 2007). Hence, if palmitoylation on cysteine residues present in Cx32-CT are critical for its ability to assemble into GJs was explored in this study. Cx32 was found to harbor three cysteine residues in its CT (Figure 1A). Results show that mutating Cx32 on single cysteine residues did not affect its trafficking or its ability to assemble into GJs in LNCaP cells (Figures 1B and 2). It was also observed that Cx32 is palmitoylated at residue C217 (Figure 4), and this palmitoylation is not required for trafficking of Cx32 or its assembly into GJs because Cx32-C217A formed GJs similar to Cx32-WT (Figure 2A). This is a novel finding showing that the site of palmitoylation in Cx32 is near the juxtamembrane position of transmembrane domain 4 at C217and that this PTM does not affect the trafficking of Cx32 or its ability to form GJs.

In contrast, the results with the triple cysteine mutant of Cx32, Cx32-3CA, showed that it did not assemble into GJs and was localized intracellularly (Figure 3). A series of double cysteine mutants were made to pinpoint if this defective phenotype was a result of mutating any two cysteines together. Results showed of all the three double mutants, only Cx32-C280A/C283A (Cx32-2CA) accumulated intracellularly, similar to Cx32-3CA (Figure 5). These data may possibly indicate that the intracellular accumulation of mutant Cx32 is due to lack of some PTM occurring at positions involving C280 and C283 or due to loss of function of a possible motif encompassing these two cysteine residues. Moreover, results also ruled out that mutating C280A and C283A together (as in Cx32-2CA), resulted in defective trafficking of Cx32 not by inhibiting palmitoylation on C217. Data showed that Cx32-2CA was palmitoylated (Figure S1), thereby suggesting that lack of palmitoylation does not result in impaired trafficking of Cx32. Findings here reveal that this defective phenotype of Cx32-2CA and Cx32-3CA is not unique to the prostate cancer cell line, LNCaP, but remain conserved across other cell lines tested, such as the pancreatic cancer cell line, BxPC3, and HEK293T cells (Figure S2). The molecular mechanism governing the intracellular accumulation of the mutants was also determined. It was found that this defective phenotype of Cx32-2CA and Cx32-3CA was due to a block in trafficking from the Golgi to the cell surface as evident from the lack of cell-surface biotinylation of these mutants (Figure 6A) and their robust colocalization with Golgi markers (Figure 6B). Moreover, intracellular accumulation was not caused by endocytosis prior to assembly (Figure 7). This highlights an important role of these cysteine residues in governing the proper transport of Cx32 through the trans-Golgi network and thus any defect will adversely affect cellular function.

Lipid PTMs like palmitoylation and prenylation are known to occur on cysteine residues. Since data here ruled out C280 and C283 to be sites of palmitoylation in Cx32, it opens up the possibility that these residues undergo some other PTM. Previous studies have identified a CAAX motif in the CT of Cx32 at position C280 (Huang et al., 2005). CAAX-motif proteins are post-translationally modified by prenylation which plays an important role in regulating protein trafficking and protein-protein interactions (Calero et al., 2003; Fukada et al., 1990). Prenylation involves the addition of a 15 or 20-carbon isoprenoid farnesylpyrophosphate (FPP) or geranylgeranylpyrophosphate (GGPP), respectively, to the cysteine residue of the target protein via a thioether bond (Zhang & Casey, 1996). The isoprenylated cysteine is part of a C-terminal CAAX box (where “C” is cysteine, “A” is usually an aliphatic residue and “X” is usually Ser, Met, Cys, Gln, Leu, Ile or Asn. The identity of this final amino acid determines if the protein becomes farnesylated (if X is Ser, Met, Cys, Gln) or geranylgeranylated (if X is Leu, Ile or Asn) (Reid et al., 2004). Interestingly, in a previous study, GenBank search revealed that Cx32 is the only cloned mammalian Cx with a CAAX-motif and that this motif is phylogenetically conserved among its orthologs. They found Cx32 to be prenylated in COS7 cells; however, this prenylation was not required for proper trafficking of Cx32 or its function in myelinating Schwann cells in mice (Huang et al., 2005).

Analysis of the amino acid sequence of Cx32 using the software PrepPS, an amino acid sequence-based predictor for various types of protein prenylation, which is based on the refinement of descriptions of sequence motifs recognized by the three enzymes [Farnesyltransferase (FT), Geranylgeranyltransferase I (GGT1), and Geranylgeranyltransferase II (GGT2)] in substrate proteins (Maurer-Stroh & Eisenhaber, 2005), indicated a high prediction score for recognition of C280 by FT (Figure S3B). The ‘X’ of the CAAX motif in Cx32 is cysteine, which favors modification by FT, as described before. Cx32 was also found to be listed in PRENbase, a database of prenylated proteins (Maurer-Stroh et al., 2007). Previous results showed that mutating Cx32 at C280 and C283 in combination (Cx32-2CA) inhibited GJ assembly (Figure 5) and these residues were critical for Cx32 trafficking to the cell surface (Figure 6). Hence it is possible that Cx32 is farnesylated at the CAAX motif and loss of this modification by mutating C280 and C283 in combination impedes its trafficking to the cell surface. To test this notion, a preliminary experiment to test if Cx32 is farnesylated in LNCaP cells have been performed. Cx32 was co-immunoprecipitated from LNCaP cells expressing Cx32-WT using an anti-Cx32 antibody and immunoblotted with an anti-farnesyl antibody. As an additional control, a similar co-immunoprecipitation assay was performed in HEK293T cells. Results showed that Cx32-WT is farnesylated in both LNCaP and HEK293T cells (Figure S3C). Collectively, these data indicate that Cx32 is prenylated. It will thus be important to analyze if this PTM of prenylation is maintained in Cx32-2CA. Absence of prenylation in this mutant will attribute its defective phenotype to lack of prenylation in Cx32 and suggest a novel role of prenylation in regulating trafficking of Cx32. Studies have shown that different types of lipidation are not mutually exclusive, in that, two or more lipids can be attached to a given protein (Aicart-Ramos et al., 2011), which would then be the case for Cx32, whereby it undergoes both palmitoylation at C217 and prenylation at C280 and C283.

Studies by Qin et.al. have shown a major role played by lysosomes in degradation of Cx43 from the early secretory compartments to the lysosome (Qin et al., 2003). Musil et al. (2000) observed that on inhibiting proteasome degradation of Cx43, GJ assembly increased and hence proposed a mechanism where gap junctional intercellular communication could be regulated at the level of Cx turnover (Musil et al., 2000). Findings here showed that the mutants Cx32-2CA and Cx32-3CA are less stable than Cx32-WT due to downregulation at the post-translational level (Figure 8) by rapid degradation in the lysosomes (Figure 9). These findings implicate that the inability of Cx32-2CA and Cx32-3CA to traffick to the plasma membrane to assemble into GJs result in enhanced degradation through the lysosomal pathway, thereby highlighting a quality control system by the cell for getting rid of mutants, detrimental to the cell.

It is well known that Cxs oligomerize as a hexamer (called connexons) in the ERGIC or in the TGN *enroute* to the cell surface (Diez et al., 1999; Musil & Goodenough, 1993); however, the underlying mechanisms regarding how connexons exit from the Golgi remain largely unknown. Studies have shown that prenylation of Ras proteins on their CAAX motif allows specific association with the ER and Golgi membranes which is required for further processing and allows engagement of a transport pathway to the PM (Choy et al., 1999). Thus, it may be possible that farnesylation of Cx32 regulates its trafficking by allowing the association of hexameric connexons with lipid rafts in Golgi membranes, and that this association directs its sorting to the cell surface. Lipid rafts, which are specialized microdomains in the PM, enriched in sphingolipids and cholesterol (Simons & Ikonen, 1997), have been proposed to be involved in protein sorting at various places in the cell (Helms & Zurzolo, 2004). A role for rafts in protein sorting in the Golgi was described for apical as well as basolateral sorting of GPI-anchored proteins in polarized epithelial cells (Brown & Rose, 1992). Alternately, this farnesylation may also regulate the hexamerization ability of the connexins to form connexons, which is required for the exit of the connexons from the Golgi.

More than 270 different mutations of the *GJB1* gene affecting different portions of Cx32 have been described for Charcot-Marie-Tooth Disease. Two of these mutations affect this potential prenylation motif: a Cys to Gly substitution at 280 (C280G) and a Ser to stop mutation at 281 (S281x) (Bergoffen et al., 1993). This underscores a possible role of lack of function of this CAAX motif in disease physiology. Thus, to summarize, the findings here show for the first time that Cx32 is palmitoylated at C217, near the juxtamembrane domain of TM4. However, this palmitoylation does not affect the trafficking of Cx32 or its ability to form GJs at the cell surface. It is known that Cxs oligomerize in the TGN (Diez et al., 1999; Musil & Goodenough, 1993), however, the underlying mechanisms regarding how connexons exit from the Golgi remain largely unknown. Data here indicates that the C280 and C283 are critical for regulating Cx trafficking, and preliminary data indicate that Cx32 is farnesylated in LNCaP cells (Figure S3). If farnesylation regulates Cx32 trafficking, then it will imply that this lipid modification of Cx32 is required for its association with lipid rafts in the Golgi, and that this association directs its sorting to the cell surface. Future studies to elucidate if farnesylation of Cx32 regulates the exit of connexons from the TGN will highlight a very important role of how this PTM affects Cx trafficking.

## MATERIALS AND METHODS

### Cell Culture

Human pancreatic cancer cell line, BxPC3 (CRL-1687), prostate cancer cell line, LNCaP(ATCC CRL 1740), and HEK293T cell (a gift from Dr. Keith Johnson, University of Nebraska Medical Center) were grown in RPMI 1640 (GIBCO) containing 5% fetal bovine serum (Sigma Aldrich) in an atmosphere of 5% CO_2_ at 37°C as described previously (Johnson et al., 2013; Katoch et al., 2015; Mitra et al., 2006). EcoPack and PTi67, the two retroviral packaging cell lines, were also obtained from ATCC and were grown as described previously (Mehta et al., 1999; Mitra et al., 2006). BxPC3 and LNCaP cells were infected with various recombinant retroviruses and pooled polyclonal cultures from approximately 2000 colonies were grown and maintained in RPMI containing G418 (200 µg/ml) (see Recombinant DNA Constructs and Retrovirus Production and Infection).

### Recombinant DNA Constructs and Retrovirus Production and Infection

The source of the retroviral vector LXSN has been described in our earlier studies (Chakraborty et al., 2010; Mitra et al., 2006). Wild-type rat Cx32 and its various mutants were cloned into pcDNA3.1 and pLXSN using PCR cloning and standard recombinant DNA protocols. Site-directed mutagenesis was used to generate mutants by QuikChange kit (Stratagene) according to manufacturer’s instructions. Recombinant DNA constructs were verified by DNA sequencing (ACGT Inc). Recombinant retroviruses were produced in EcoPack and PTi67 packaging cell lines. BxPC3 and LNCaP cells were multiply (2-4 times) infected with various recombinant retroviruses produced from PTi67 cells and selected in G418 (400 μg/ml) as described (Chakraborty et al., 2010; Govindarajan et al., 2010; Mehta et al., 1999). Pooled cultures from about 2000 colonies obtained from 2-4 dishes were expanded, frozen, and maintained in selection media containing G418 (200 µg/ml). Pooled polyclonal cultures were used within 3-5 passages for immunocytochemical and biochemical analyses.

### Antibodies and Immunostaining

A mouse hybridoma M12.13 secreting monoclonal antibody against rat Cx32 (Goodenough et al., 1988) was a gift from Dr. Dan Goodenough (Harvard University). This monoclonal antibody recognizes residues 111-125 of rat, mouse and human Cx32. In some experiments, we also used two rabbit polyclonal antibodies raised against residues 106-124 and 262-279 of carboxyl tail of Cx32 (Sigma; C-3595 and C-3470). Other rabbit polyclonal antibodies used in this study are as follows: anti-GM130 (610822, BD Transduction Laboratory); anti-TGN46 (ab5095, Abcam); anti-β-catenin (C2206) and farnesyl antibody (AB4073) from Sigma. Other mouse monoclonal antibodies used in this study are as follows: anti-E-cadherin, anti-β-catenin and anti-α-catenin (gift from Dr. Keith R. Johnson, University of Nebraska Medical Center). For immunostaining 3 x10^5^ BxPC3 and 3 x10^5^ LNCaP cells were seeded on glass cover slips in six-well clusters and allowed to grow for 72 h, after which they were fixed with 2% paraformaldehyde and immunostained as described previously (Chakraborty et al., 2010; Govindarajan et al., 2010; Johnson et al., 2013; Katoch et al., 2015). Anti-rabbit and anti-mouse secondary antibodies, conjugated with Alexa 488 or Alexa 594 (Invitrogen), were used as appropriate.

### Fluorescence Microscopy

Fluorescent images of immunostained cells were acquired with a Leica DMRIE microscope (Leica Microsystems, Wetzler, Germany) using a 63x oil objective (NA 1.35). This microscope is also equipped with a Hamamatsu ORCA-ER2 CCD camera (Hamamatsu City, Japan). Co-localization was measured in z-stacked images taken 0.3 µm apart using the commercial image analysis program Volocity 6.0.1 (Improvision, Lexington, MA) as described previously (Chakraborty et al., 2010; Govindarajan et al., 2010; Johnson et al., 2013; Katoch et al., 2015). Saturation of the detector, which would alter background adjustment, was prevented by minimizing exposure of each fluorescently-tagged antibody during acquisition. A background-corrected Pearson’s correlation coefficient was used to determine fluorescence co-localization as described (Barlow et al., 2010). Images were captured with a Z-plane thickness of 0.5 uM and processed with the Zen Blue software (Carl Zeiss). The processed images were imported into Volocity, cropped as appropriate, and exported as TIFF files.

### Gap Junction Size Measurement

The surface area of a GJ plaque was measured as follows. After acquiring images of immunostained cells, serial *z*-sections (0.3 µm) were collected and subjected to iterative volume deconvolution using Volocity image-processing software (Improvision). The de-convolved images of single optical sections were used for measuring GJ size. Each distinct fluorescent punctum at the cell-cell contact site, delineated by E-cadherin immunostaining, was considered as one GJ plaque. The area of each GJ plaque was calculated by drawing an ROI around each punctum using the free hand ROI tool of the “Measurement” module of Volocity. The area of each GJ is represented as “pixel count” where one pixel corresponded to 0.01 μm^2^. In each captured image, 3-5 puncta were randomly chosen for the measurement of area and 5-15 images were used. We used the “Extended focus” display option, which merges all captured z planes, for determining the number of GJ plaques per cell-cell interface. The number of distinct visible puncta along the cell-cell interfaces of two adjoining cells were then counted. We chose only distinct interfaces at random to avoid ambiguity in localization of the puncta. Typically, 60-70 interfaces were measured from three independent experiments.

### Detergent Extraction and Western Blot Analysis

BxPC3 (3×10^6^) and LNCaP (2×10^6^) cells, seeded in replicate 10 cm dishes in 10 ml of complete medium, were grown for 72 h. Cell lysis, detergent-solubility in 1% Triton X-100 (TX100), and Western blot analysis were performed as described previously (Chakraborty et al., 2010; Govindarajan et al., 2010; Katoch et al., 2015; Mitra et al., 2006). Normalization was based on equal cell number for the analysis of detergent-soluble and -insoluble fractions by SDS-PAGE analysis of cell lysates.

### Cell Surface Biotinylation Assay

For cell surface biotinylation, BxPC3 (5×10^5^) and LNCaP (4×10^5^) cells expressing Cx32-WT and various mutants were seeded in 6 cm dishes in replicates and grown to 70-80% confluence. Biotinylation reaction was carried out at 4°C for 1 h with freshly prepared EZ-LinkSulfo-NHS-SS Biotin reagent (Pierce) at 0.5 mg/ml in phosphate buffered saline (PBS) supplemented with 1 mM CaCl_2_ and 1 mM MgCl_2_. The affinity precipitation of biotinylated proteins was from 200 μg of total protein using 100 µl of streptavidin-agarose beads (Pierce) on a rotator overnight at 4°C. SDS-PAGE followed by Western blotting was used to resolve the streptavidin-bound biotinylated proteins after elution as described previously (Chakraborty et al., 2010; Govindarajan et al., 2010; Govindarajan et al., 2002). The kinetics of degradation of cell-surface-associated Cx32-WT and mutants was determined as follows. After biotinylation, washing and quenching biotin, biotinylated proteins were chased at 37°C for various times before affinity precipitation with streptavidin as described previously (Govindarajan et al., 2010; Johnson et al., 2013; Katoch et al., 2015). The protein concentration was determined using the BCA reagent (Pierce).

### Immunoprecipitation

For co-immunoprecipitation experiments, HEK293T cells were grown in 6-cm dishes to 80% confluence and transfected with 5 µg of each plasmid for cotransfection studies. Twenty-four hours post-transfection, the cells were harvested and lysed in a non-denaturing Tris-NP40 lysis buffer (10 mM Tris-HCl, pH 8.0, 0.5% NP-40, 1 mM EDTA) supplemented with 1 mM PMSF, 2 mM Na_3_VO_4_, 1 mM NaF, and a protease inhibitor cocktail. Following lysis and protein estimation, 500 µg of total protein was incubated with antibodies and mixed overnight at 4 °C. Next day, anti-mouse beads (Sigma) was washed with Tris-NP40 buffer once. The cell extract-antibody mix was incubated with IgG affinity gel and mixed at 4 °C for 2 h. The immune complexes were washed 4 times with the wash buffer (10 mM Tris-HCl, pH 7.5, 150 mM NaCl, and 0.25% Tween 20). After the final wash, the beads were suspended in 2X Laemmli sample buffer and incubated at 37°C for 45 min to elute bound proteins. Proteins were then analyzed by SDS-PAGE and Western blotting.

### Cell Transfection

BxPC3 (4×10^5^) and LNCaP (5 x10^5^) cells were seeded on glass cover slips in 6-well clusters 24 h prior to transfection. Cells were transfected with various plasmids in duplicate using XtremeGENE (Roche Diagnostics) according to the manufacturer’s instructions. Two μg of plasmid DNA per well was used for transfection. When cells were to be co-transfected with two plasmids, one µg of each plasmid was used. Expression was analyzed 24 h post-transfection after fixing and immunostaining cells with the desired antibodies as described previously (Govindarajan et al., 2010; Johnson et al., 2013; Katoch et al., 2015; Mitra et al., 2006). For transfection of HEK293T for immunoprecipitation studies, replicate 6 cm dishes were transfected with 2.5 µg of two plasmids and cells were lysed and immunoprecipitated as described above.

### Hypertonic Sucrose Treatment and K^+^ depletion

For hypertonic sucrose and K^+^ -depletion treatments, 3×10^5^ LNCaP cells expressing Cx32-WT and the Cx32-cysteine mutants were seeded on glass cover slips in six-well clusters and allowed to grow to confluence for 72 h. Cells were incubated for 1 h at 37^0^C in starvation medium (RPMI, 0.5 % BSA, 30 mM HEPES, pH 7.5). Cells to be subjected to K^+^ depletion were pre-treated as follows: Cells were rinsed 3 times with K^+^-free buffer (20 mM HEPES, pH, 140 mM NaCl, 1 mM CaCl_2_, 1 mM MgCl2, mM D-glucose), followed by incubation for 5 min in hypotonic K^+^-free buffer (K^+^-free buffer diluted 1:1 with water), washed again 3 times with K^+^-free buffer, and incubated in it for 15 min at 37 ^0^C. Coverslips were then transferred to chilled binding medium (starvation medium containing Alexa 594-conjugated with Transferrin at 25 µg/ml) and incubated for 1 h at 4 ^0^C. Cells were then washed 3 times with chilled PBS. Cells to be subjected to hypertonic sucrose treatments were incubated in hypertonic sucrose medium (0.45 M sucrose) for 2 h and cells to be subjected to K^+^ -depletion were incubated in warm K^+^-free buffer for 2 h. Control cells were incubated in starvation medium for 2 h. At the end of the treatments, cells were fixed and immunostained for Cx32 as described above.

### Treatments

Stock solutions of various reagents were prepared as follows: Leupeptin at 100 mM in water; NH4Cl at 1 M in water (SIGMA); Lactacystin at 20 mM in ethanol; brefeldin at 10 mM in DMSO (BIOMOL), Cycloheximide at 1 mg/ml in DMSO (Sigma). These solutions were stored at -20 ^0^C in small aliquots. All solutions were appropriately diluted in the cell culture medium at the time of treatment. Controls were vehicle treated.

### ABE Pulldown Assay

The Acyl Biotin Exchange (ABE) assay was carried out as described (Wan et al., 2007) with a few modifications. Briefly, cells were grown in six 100 mm dishes to confluency. They were then washed with cold PBS. Lysis buffer (Wang et al.) (50 mM Tris-HCl, pH 7.4, 5 mM EDTA, 150 mM NaCl) containing 10 mM NEM was added to the dishes to block free thiols. Cells were then harvested, lysed and sonicated. Triton X-100 was then added to a final concentration of 1.7% and incubated with gentle end-over-end rotation at 4°C for 1 hr. Each cell lysate was collected by centrifugation at 14000 rpm for 15 min at 4°C. The proteins were then precipitated with chloroform/methanol (CM) (2:5 vol/vol) and resuspended in SB buffer (4% SDS, 50 mM Tris-HCl, pH 7.4, 5mM EDTA) with 10mM NEM and then incubated at 37°C with gentle agitation for 2-3 min until it became clear. LB containing 1 mM NEM, 0.2 % TX100, protease inhibitor cocktail (PIC), and phenylmethylsulfonylfluoride (PMSF) was added to each sample and the solution was incubated overnight at 4°C with gentle mixing. NEM was removed from the samples by three sequential CM precipitations, and the samples were finally resuspended in 300 μl SB. The samples were then divided into two equal portions. One portion was incubated with HA buffer (1 M hydroxylamine, pH 7.4, 150 mM NaCl, 0.2 % TX100, PIC, 1 mM PMSF, 1 mM HPDP-biotin) and the other with the same buffer minus HPDP-biotin, as a control, at room temperature for 1 h with gentle mixing. Then the proteins were CM precipitated once to remove unreacted HPDP-biotin and resuspended in SB and incubated for 1 hr. This was followed by three sequential CMs. Finally, the protein was dissolved in LB with 0.2 % TX-100 and incubated with gentle end-over-end rotation at 37°C for 30 min. This was followed by centrifugation at 13000 rpm for 5 min. The supernatant was collected and used for protein estimation. 200 ug of the protein was incubated with 30 µl of streptavidin agarose beads and mixed with gentle end-over-end rotation at 37°C for 90 min. The beads were then washed 4 times in cold 1X TBST (10 mM Tris-HCl pH 7.5, 150 mM NaCl and 0.25% Tween 20). After the final wash, the beads were suspended in 2X Laemmli sample buffer and incubated at 37°C for 45 min to elute bound proteins. Proteins were then analyzed by SDS-PAGE and Western blotting

### mRNA isolation and quantitative Real-Time PCR

Total RNA from cells was isolated using RNeasy mini kit (Qiagen, 74104) according to manufacturer’s protocol. cDNA was prepared from 1μg of total RNA using Transcriptor First Strand Synthesis Kit (Roche, 04379012001) with random hexamer primer. Quantitative real-time PCR (qRT-PCR) was carried out with Power SYBR Green master mix (Life Technologies, 4368706) in an Applied Biosystems machine using primers specific to Cx32 and 36B4 (Acidic Ribosomal Phosphoprotein P0). Each qRT-PCR experiment was carried out in duplicate and average cycle thresholds (Cts) were obtained for each sample. Briefly, the Ct of the reference gene was subtracted from the Ct value of the target gene to obtain delta (δ) Ct values (δCt). Then the δCt of each of the samples, from the was compared with the average δCt from the WT sample to get δδCt values. The fold change in expression between the samples was calculated as 2^-δδCt^.

### Statistical Analysis

Data analysis was done using GraphPad Prism version 8.0. (Comparisons between two groups were made using Student’s unpaired two-tailed *t* test. Data are expressed as mean ± SEM (standard error of mean). Values of p < 0.05 were considered to be statistically significant.

## ACKNOWLEDGEMENTS

I express my sincere gratitude to Dr. Parmender P. Mehta, University of Nebraska Medical Center, under whose supervision the above work was carried out. In his absence, owing to his unfortunate demise, I am taking this work forward. I thank Dr. Samikshan Dutta for critically reviewing this manuscript and providing key suggestions. I thank Dr. Keith Johnson, University of Nebraska Medical Center, and his lab members for help with the ABE assay. Finally, I thank my current postdoctoral supervisor, Dr. Timothy McGraw, Weill Cornell Medicine, for his time and effort in providing critical insights into this manuscript.

---

**Figure S1.**
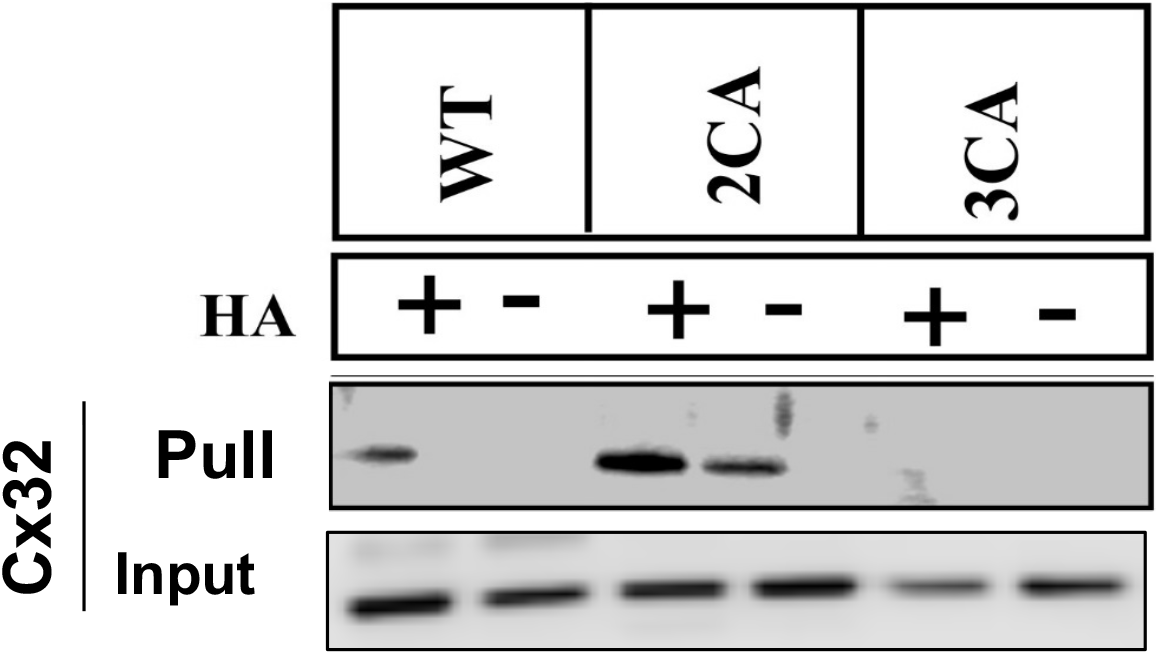
Cx32-2CA is palmitoylated. ABE Pulldown assay. Lysates from LNCaP cells expressing Cx32-WT, Cx32-2CA or Cx32-3CA were split into two halves and treated with HA (+) or left untreated (-). After the streptavidin-agarose pull-down, the captured proteins were resolved by SDS-PAGE and immunoblotted for Cx32 and p120. (See Materials and Methods). For the input, 10 μg of total protein was used and immunoblotted for Cx32. Note that Cx32-2CA is palmitoylated, similar to Cx32-WT.

**Figure S2.**
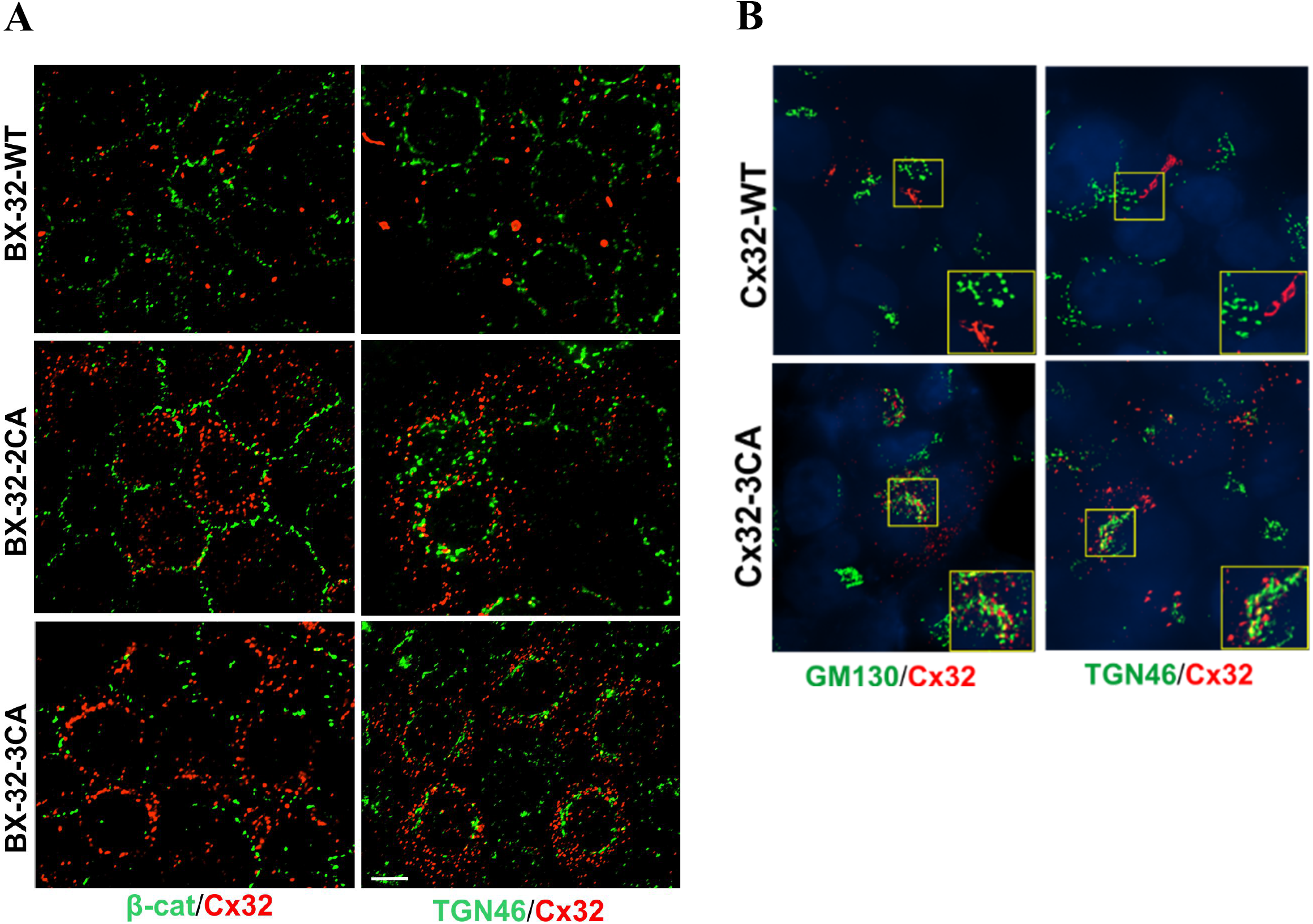
Cx32-2CA and Cx32-3CA accumulates intracellularly in BxPC3 and HEK293T cells. **A**. BxPC3 cells retrovirally expressing Cx32-WT, Cx32-2CA or Cx32-3CA were immunostained for Cx32 (red) and β-catenin (β-cat, a cell-surface marker) and TGN46, a Golgi. marker (green). Note that the mutants showed an intracellular localization. **B**. HEK293T cells transiently expressing Cx32-WT, Cx32-2CA or Cx32-3CA were immunostained for Cx32 (red) and Golgi markers, GM130 and TGN46 (green). Note that the mutants showed an intracellular localization.

**Figure S3:**
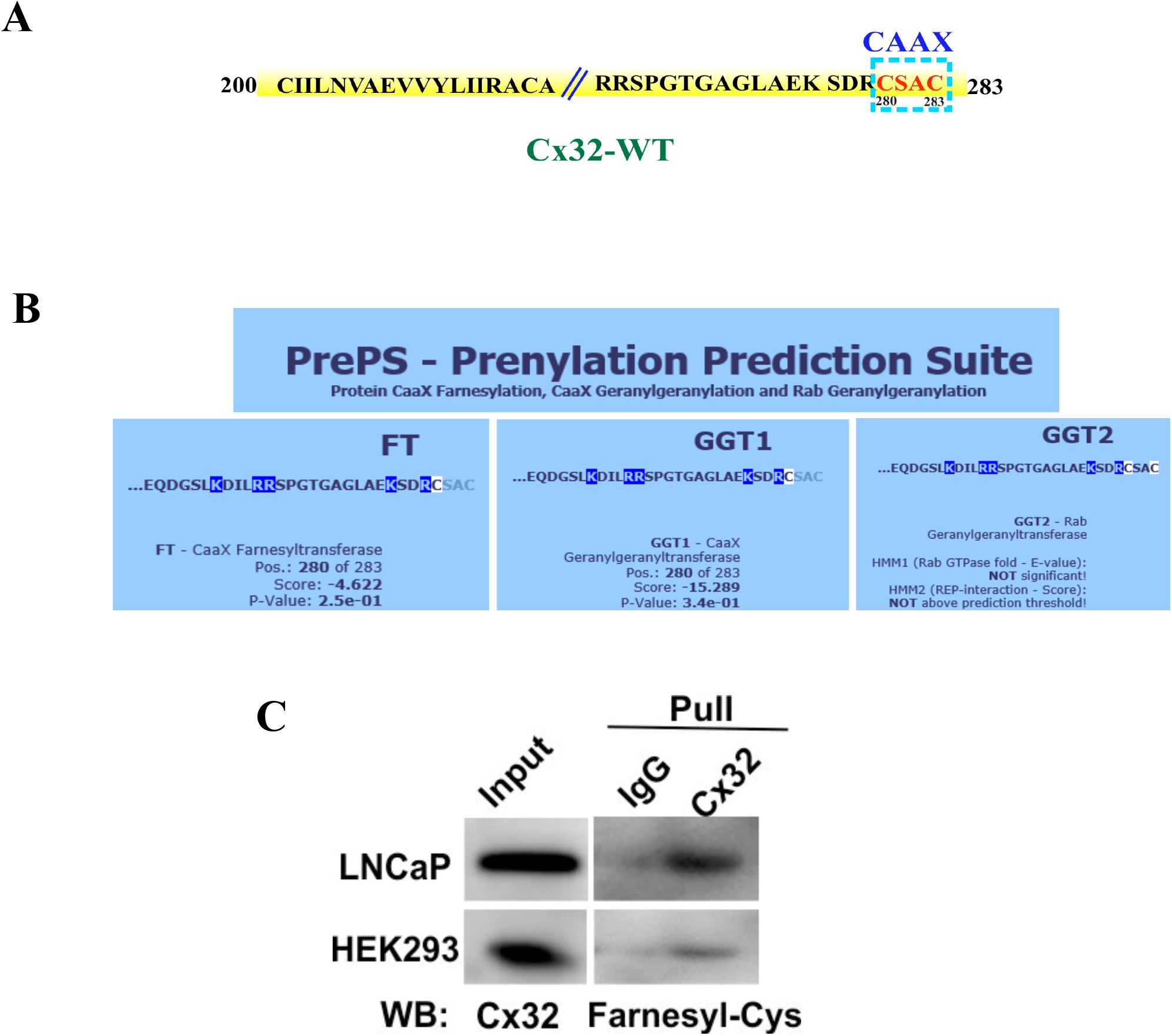
Cx32-WT is farnesylated. **A**. The CAAX motif for Cx32 is located at position C280 of its CT, circumscribed within a blue rectangle. **B**. A snapshot of the prediction obtained using the PrePS software. Note that as compared to GGT1 (Geranylgeranyltransferase I) and GGT2 (Geranylgeranyltransferase II), the prediction score of FT (Farnesyltransferase) acting on position C280 is the highest. **C**. Whole cell lysates from LNCaP cells and HEK293T cells expressing Cx32-WT were pulled down using an anti-Cx32 antibody and immunoblotted with an anti-farnesyl antibody. For the input, 10 μg of total protein was used and immunoblotted for Cx32.

